# The transcription factor Rreb1 regulates epithelial architecture and invasiveness in gastrulating mouse embryos

**DOI:** 10.1101/2020.11.12.379578

**Authors:** Sophie M. Morgani, Jie Su, Jennifer Nichols, Joan Massagué, Anna-Katerina Hadjantonakis

**Author notes:** Corresponding authors: Sophie M. Morgani, PhD and Anna-Katerina Hadjantonakis, PhD.

## Abstract

Ras-responsive element-binding protein 1 (Rreb1) is a zinc-finger transcription factor downstream of RAS signaling. *Rreb1* has been implicated in cancer but little is known about its role in mammalian non-disease states. Here, we found that Rreb1 is essential for mouse embryonic development. Loss of *Rreb1* led to a reduction in the expression of vasculogenesis factors, cardiovascular defects and embryonic lethality. During gastrulation, the absence of *Rreb1* also resulted in the upregulation of cytoskeleton-associated genes, a change in the organization of F-ACTIN and adherens junctions within the pluripotent epiblast, and perturbed epithelial architecture characterized by irregular tissue folding and abnormal accumulations of cells. Moreover, *Rreb1* mutant cells ectopically exited the epiblast epithelium through the underlying basement membrane, paralleling cell behaviors observed during metastasis. Thus, disentangling the function of Rreb1 in development could shed light on its role in cancer and other diseases involving loss of epithelial integrity.

## 1. Introduction

Ras-responsive element-binding protein 1 (RREB1) is a zinc-finger transcription factor that acts downstream of RAS (Thiagalingam et al., 1996). It is evolutionarily conserved (Ming, Wilk, Reed, & Lipshitz, 2013), widely-expressed (FujimotoNishiyama, Ishii, Matsuda, Inoue, & Yamamoto, 1997), can function both as a transcriptional repressor and activator (Deng, Xia, Zhang, Ejaz, & Liang, 2020), and interacts with several signaling pathways, including EGFR/MAPK (M. Kim et al., 2020) and JNK/MAPK (Melani, Simpson, Brugge, & Montell, 2008; Reed, Wilk, & Lipshitz, 2001), which regulate RREB1 through phosphorylation, and JAK/STAT (Melani et al., 2008), TGF-β/SMAD (Su et al., 2020), Notch, and Sonic Hedgehog (J. J. Sun & Deng, 2007), which cooperate with RREB1 in transcriptional regulation. These properties suggest that RREB1 plays key contextual biological roles.

Most of what we know about mammalian Rreb1 stems from cancer studies where mutations in, or altered expression of, this gene have been associated with leukemia (Yao et al., 2019), melanoma (Ferrara & De Vanna, 2016), thyroid (Thiagalingam et al., 1996), and prostate (Mukhopadhyay et al., 2007) cancers, as well as pancreatic and colorectal cancer metastasis (Cancer Genome Atlas Research Network. Electronic address & Cancer Genome Atlas Research, 2017; Hui et al., 2019; Kent, Sandi, Burston, Brown, & Rottapel, 2017; Li et al., 2018). However, the function of mammalian Rreb1 in normal, non-disease states remains unclear.

The Drosophila homolog of Rreb1, *Hindsight* (hnt, also known as *pebbled*), is required for embryonic development (Wieschaus, Nussleinvolhard, & Jurgens, 1984) where it regulates cell-cell adhesion and collective migration in various contexts, including trachea and retinal formation, border cell migration, and germ-band retraction (Melani et al., 2008; Pickup, Lamka, Sun, Yip, & Lipshitz, 2002; Wilk, Reed, Tepass, & Lipshitz, 2000). Additionally, we recently reported that chimeric mouse embryos containing *Rreb1* mutant cells exhibit early embryonic phenotypes (Su et al., 2020), indicating that *Rreb1* has a role in mammalian development. The fundamental biological processes that regulate development are frequently hijacked in cancer. A notable example is the specification and patterning of the embryonic germ layers, known as gastrulation. Gastrulation involves an epithelial-mesenchymal transition (EMT), basement membrane remodeling, and collective cell migration, processes that also coordinately drive cancer progression (Aiello & Stanger, 2016; Cofre & Abdelhay, 2017). Thus, characterizing the mechanisms and identifying critical factors that control development will shed light on how they are dysregulated in disease.

Here we generated a *Rreb1* mutant mouse line and investigated its role and requirement during mouse embryonic development. We found that *Rreb1* is expressed within both the embryo-proper and the extraembryonic supporting tissues and is essential for a variety of processes including neural tube closure and vasculogenesis. Loss of *Rreb1* resulted in a change in the organization of the cytoskeleton and adherens junctions, increasingly variable cell orientation, irregular folding, and the emergence of aberrant cell masses within the pluripotent epiblast epithelium during gastrulation. Furthermore, a fraction of *Rreb1*^−/−^ epiblast cells breached the underlying basement membrane, and ectopically exited the epithelium, seeding epiblast-like cells throughout the embryo. These data collectively demonstrated that *Rreb1* is required to maintain epithelial architecture during mammalian development and loss of this factor promotes cell behaviors reminiscent of those observed in metastasis. Thus, future studies to unravel the tissue-specific targets and mechanism of action of *Rreb1* during development may also shed light on its role in disease states.

## 2. Results

### *Rreb1* is expressed as cells exit pluripotency

We first characterized the pattern of expression of *Rreb1* in the early mouse embryo using single-cell transcriptomic (scRNA-seq) datasets, previously generated by us and others (Nowotschin et al., 2019; Pijuan-Sala et al., 2019). These revealed that, at pre-implantation and early post-implantation stages (embryonic day (E) 3.5-5.5), *Rreb1* is expressed by trophectoderm cells that form the fetal portion of the placenta, the inner cell mass (ICM) that gives rise to embryonic epiblast and extraembryonic primitive endoderm, and its descendants contributing to the yolk sac (Figure S1A) (Nowotschin et al., 2019). At the onset of gastrulation (E6.5), *Rreb1* is expressed within trophectoderm-derived extraembryonic ectoderm (ExE), primitive endoderm-derived visceral endoderm (VE), and the epiblast-derived primitive streak, where cells undergo an EMT and start to differentiate into the mesoderm and endoderm germ layers (Figure S1A) (Pijuan-Sala et al., 2019). From E7.75 onwards*, Rreb1* is broadly expressed including in the epiblast, primitive streak, neurectoderm, mesoderm, and definitive endoderm (DE) (Figure S1A) (Pijuan-Sala et al., 2019).

We validated these scRNA-seq data in wholemount preparations of embryos expressing a LacZ-tagged transcriptional reporter (S1B Fig, European Conditional Mouse Mutagenesis Program) (Bradley et al., 2012), and confirmed *Rreb1*^LacZ^ expression within the ICM and trophectoderm of the blastocyst (Figure S1C), as well as within the VE before gastrulation (E5.5, Figure 1A), the VE, primitive streak, embryonic and extraembryonic mesoderm (cells derived from the primitive streak), and distal anterior epiblast during gastrulation (E6.5-7.5, Figure 1A, S1D), and within the yolk sac endoderm, node, notochord, primitive streak, blood, allantois, head mesenchyme, and pharyngeal arches at E8.0-10.5, around midgestation (Figure 1A, S1D-H). A comparable expression pattern was detected by mRNA *in situ* hybridization (Figure S1I). At E10.5, *Rreb1*^LacZ^ was expressed in regions of high FGF signaling activity (Morgani, Saiz, et al., 2018), including the limb buds, frontonasal processes and isthmus (Figure S1G). Furthermore, the expression domain of *Rreb1*^LacZ^ within the tail bud varied between individual embryos (Figure S1G), suggesting that it is transcriptionally regulated by the segmentation clock. *In vitro*, *Rreb1*^LacZ^ marked a subpopulation of pluripotent embryonic stem cells and epiblast stem cells, when maintained under self-renewing conditions, and became more widely expressed as cells were driven to differentiate by the removal of the cytokine LIF or addition of FGF (Figure 1B). Thus, *Rreb1* is expressed in the embryonic lineages as pluripotency is exited and the germ layers are specified, and in the extraembryonic tissues.

**Figure 1.**
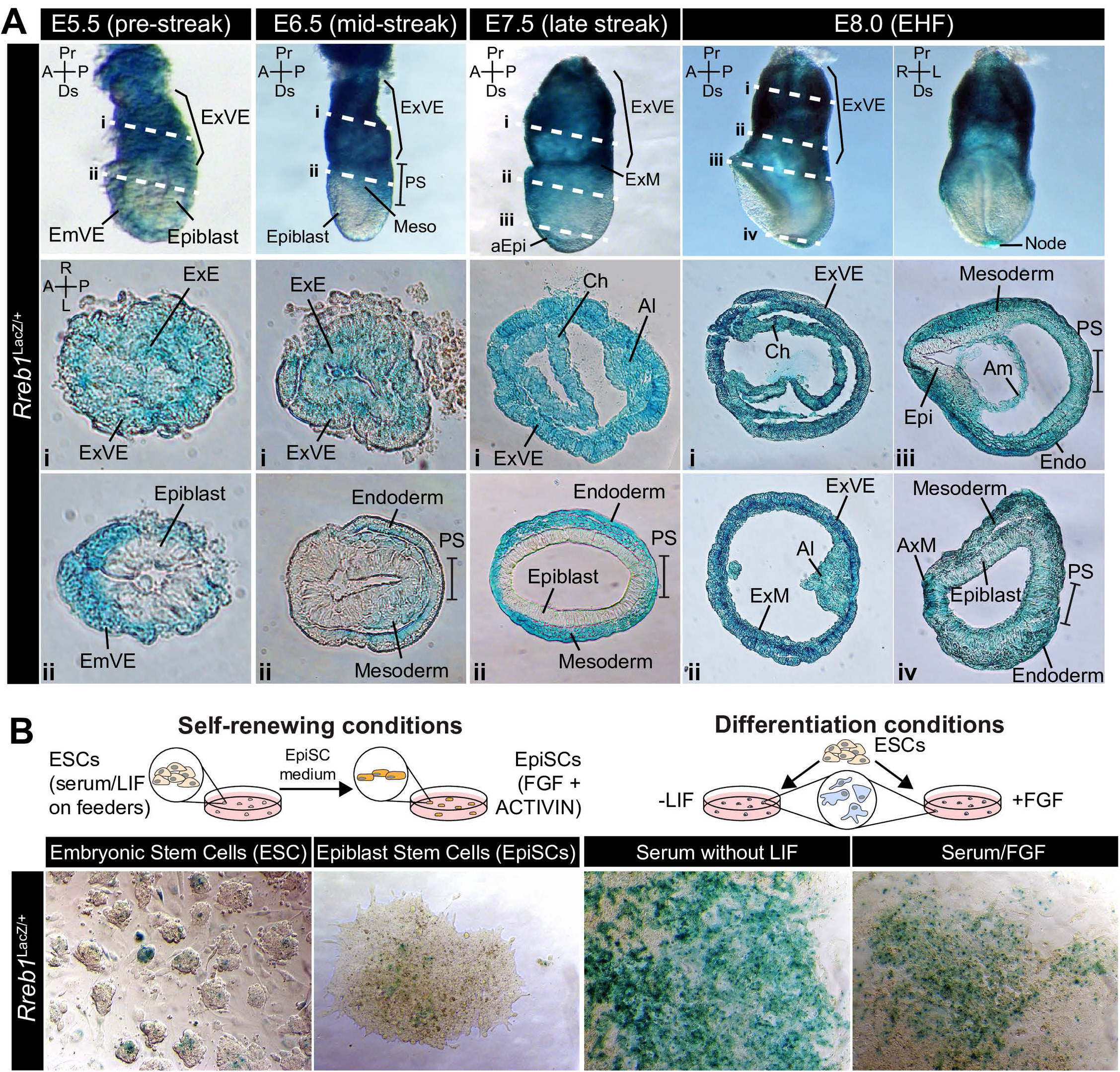
*Rreb1* is expressed within embryonic and extraembryonic tissues. **A.** Wholemount images of *Rreb1*^LacZ/+^ mouse embryos from embryonic day (E) 5.5-8.5. Dashed lines mark approximate plane of transverse sections shown in lower panels. Section iii from E7.5 is located in Figure S1D. **B.** *Rreb1*^LacZ^ reporter mouse embryonic stem cells (mESCs) (i) and epiblast stem cells (ii) under self-renewing conditions. mESCs were grown in serum/LIF on feeders. Panels (iii) and (iv) show mESCs after 7 days of differentiation in the absence of LIF or in the absence of LIF plus 12 ng/ml FGF2. A, anterior; P, posterior; Pr, proximal; Ds, distal; L, left; R, right; ExM, extraembryonic mesoderm; ExVE, extraembryonic visceral endoderm; AVE, anterior visceral endoderm; aEpi, anterior epiblast; Meso, mesoderm; Endo, endoderm; Epi, epiblast; PS, primitive streak; Am, amnion; Al, allantois; Ch, chorion; AxM, axial mesoderm.

### *Rreb1* is essential for mouse embryonic development

We previously used *Rreb1*^−/−^ cells to generate chimeric mouse embryos and found that these exhibited severe morphological defects during gastrulation (Su et al., 2020). To interrogate the developmental function of Rreb1, we proceeded to generate a *Rreb1* knockout mouse using CRISPR-Cas9 technology (Figure 2A, Materials and methods). *Rreb1*^+/−^ mice were viable and fertile, but heterozygous intercrosses yielded no homozygous mutant offspring. From E7.5 onwards, mutant embryos were smaller than wild-type littermates (Figure 2B, C, S2A, B) and, based on morphology and somite number, were approximately 9.5 hours retarded (Figure S2B, C). At E9.0-9.5, *Rreb1*^−/−^ embryos exhibited various defects, including microcephaly (2/7 at E9.5, Figure S2D), an open foregut (Figure S2E), and an open neural tube at the forebrain, midbrain, and posterior neuropore level (8/10 *Rreb1*^−/−^ at E9.5, Figure 2D-E, S2F). *Rreb1*^−/−^ embryos with open neural tubes were recovered, albeit at lower frequencies, at E10.5 (2/8 *Rreb1*^−/−^, Figure 2B, 3F), indicating that this phenotype is partially associated with the developmental delay.

**Figure 2.**
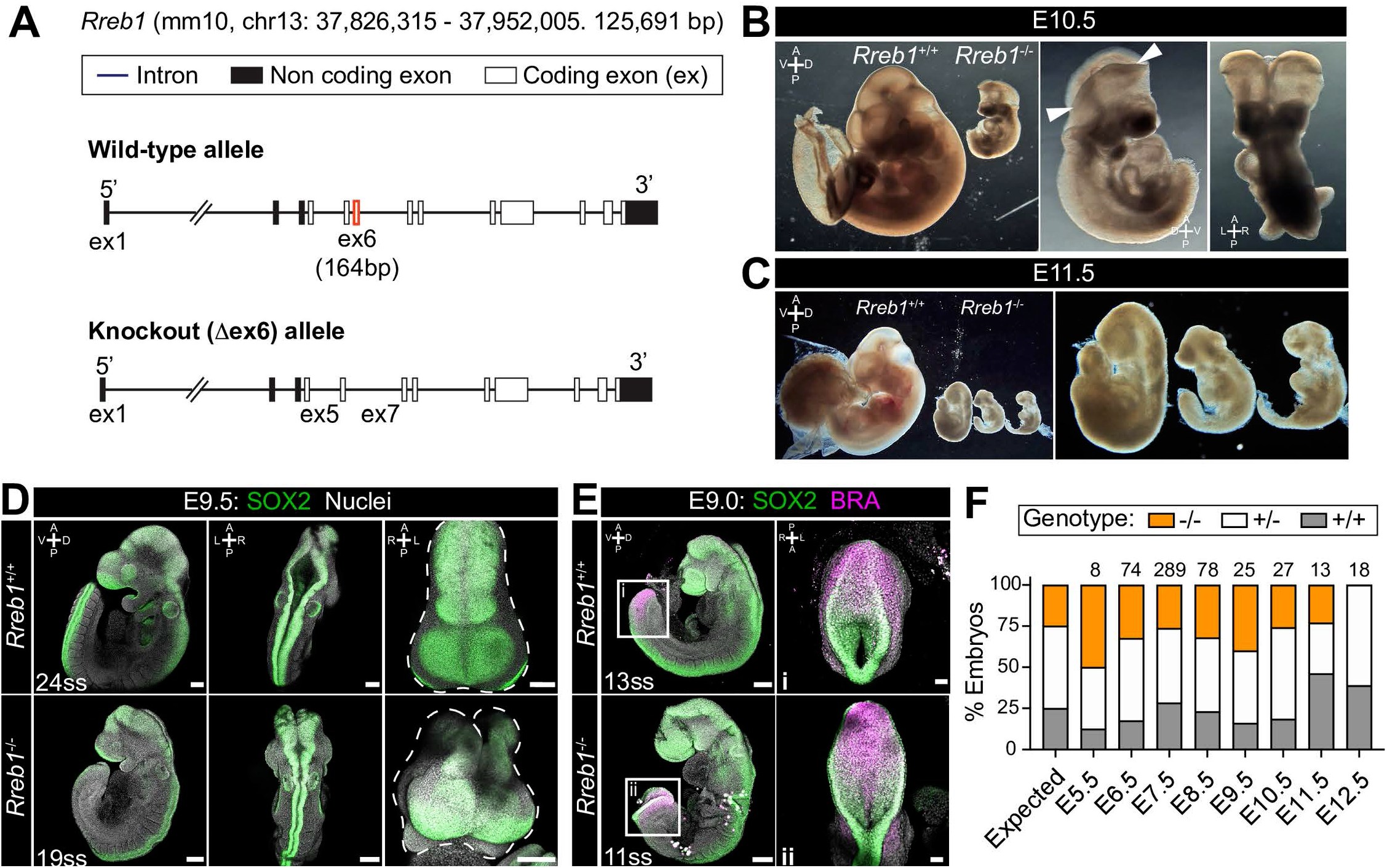
*Rreb1* is necessary for mouse embryonic development. **A.** Schematic diagram showing the strategy used to generate the *Rreb1* mutant allele. CRISPR-Cas9 was used to delete the majority of the coding DNA sequence of Exon 6. We created a large (approximately 700 bp) and small (approximately 540 bp) deletions. Both lines exhibited comparable phenotypes, thus we combined these data. UTR, untranslated region. **B-C.** Brightfield images of *Rreb1*^+/+^ and *Rreb1*^−/−^ littermates at E10.5 and E11.5 Arrowheads indicate boundary of open neural tube. Righthand panels show mutant embryos at higher magnification. **D-E.** Confocal maximum intensity projection (MIP) of wholemount E9.0 and 9.5 mouse embryos, sb 200 μm. Somite pair numbers (ss) shown on the images. **D.** Right panel shows a MIP frontal view and outline (dashed line) of the head of the embryo emphasizing the neural tube closure defects in the *Rreb1*^−/−^. **E.** Box highlights image of posterior neuropore shown in high magnification in adjacent panel, sb 100 μm. **F.** Bar chart summarizing the percentage of *Rreb1*^+/+^, *Rreb1*^+/−^ and *Rreb1*^−/−^ embryos recovered at each developmental stage. The first bar indicates the expected Mendelian ratios of each genotype. N numbers are shown above each bar. D, dorsal; V, ventral; A, anterior; P, posterior; L, left; R, right; VE, visceral endoderm; ExE, extraembryonic ectoderm; DE, definitive endoderm; PS, primitive streak; Epi, epiblast; Meso, mesoderm; fb, forebrain; mb, midbrain; hb, hindbrain; ys, yolk sac.

Additionally, mutant embryos displayed aberrant notochord formation. In wild-type embryos, the axial mesoderm, marked by BRACHYURY expression in cells anterior to the gut tube, gives rise to the prechordal plate rostrally (Figure S2G i) and to the tube-like notochord caudally (Figure S2G ii-iv) (Balmer, Nowotschin, & Hadjantonakis, 2016). However, in *Rreb1*^−/−^, BRACHYURY-expressing cells did not establish a tube, instead, intercalating into the foregut (Figure S2G v), protruding into the foregut lumen (Figure S2G vi), or generating multiple distinct clusters (Figure S2G vii). Thus, loss of *Rreb1* results in a range of phenotypic abnormalities initiating at gastrulation and resulting in midgestation lethality.

Homozygous mutants began to be resorbed at E11.5, indicated by the disintegration of embryonic tissues (Figure 2C), and were not recovered at E12.5 (Figure 2F). Thus, *Rreb1* is essential for mouse development, where it regulates a variety of processes.

### *Rreb1* is required for cardiovascular development

Rreb1 is a transcription factor that functions as a context-dependent repressor or activator (Deng et al., 2020). To define the transcriptional changes associated with a developmental loss of *Rreb1* and gain insights into its mechanism of action, we performed RNA-sequencing of *Rreb1*^−/−^ embryos and compared them to wild-type (*Rreb1*^+/+^) transcriptomes. Embryos were isolated and analyzed at E7.5 (Figure 3A), coinciding with the emergence of overt morphological defects resulting from loss of *Rreb1* (Figure S2A, B). We identified 65 genes that were significantly downregulated and 200 that were upregulated in *Rreb1*^−/−^ vs. *Rreb1*^+/+^ embryos (fold-change >log2(2), *p* <0.05, Table S1).

**Figure 3.**
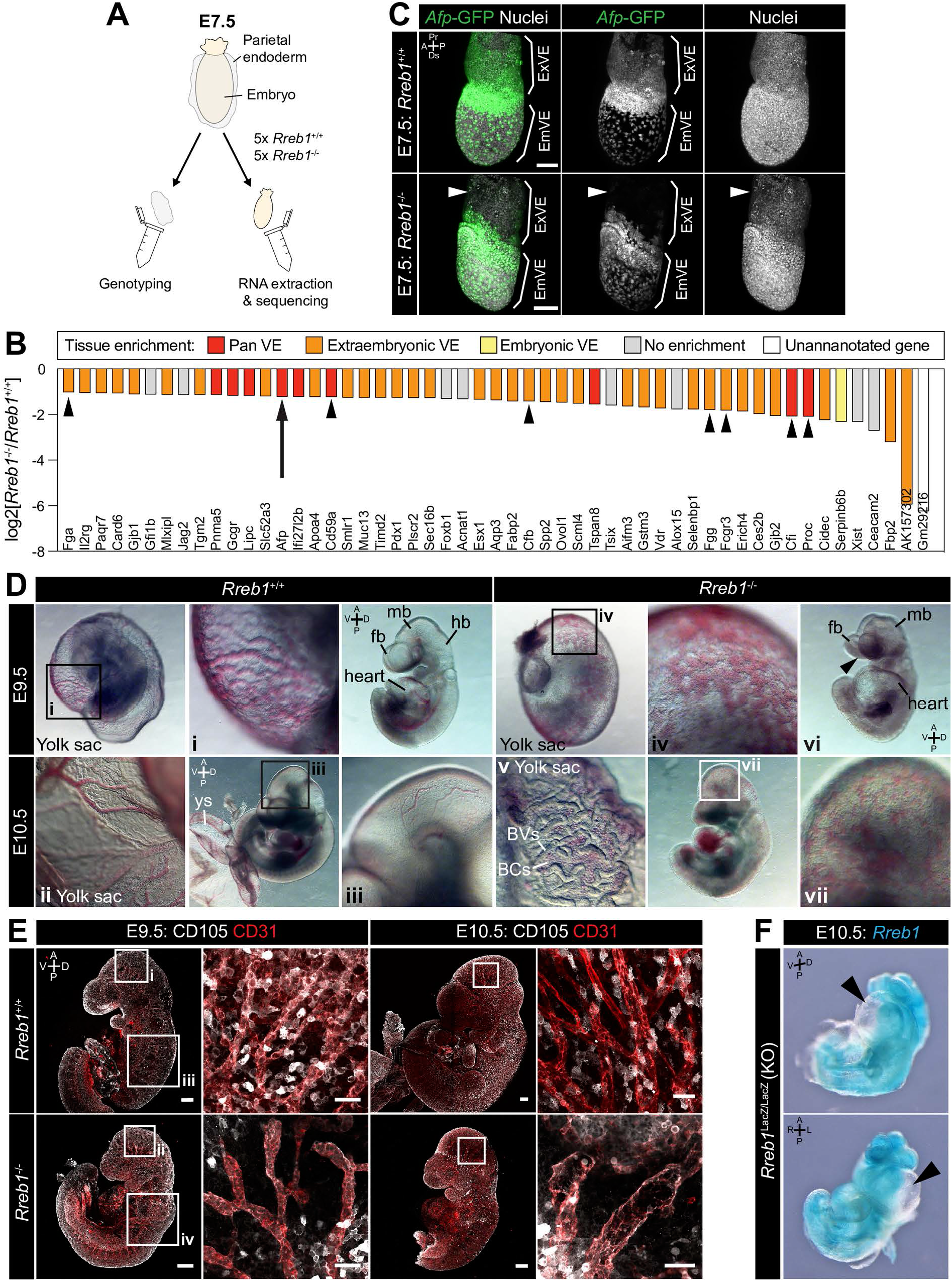
Loss of *Rreb1* causes cardiovascular defects in the early mouse embryo. **A.** Schematic diagram depicting the sample collection methodology for whole embryo RNA-seq. Individual embryos were isolated from the uterus and the parietal endoderm dissected, lysed, and used for genotyping. The remaining part of the embryo was used for RNA extraction. Following genotyping, 5 individual wild-type and 5 individual mutant embryos were selected for sequencing. **B.** Graph showing the list of significantly downregulated genes in *Rreb1*^−/−^ versus *Rreb1*^+/+^ embryos that were detected via single-cell sequencing in (Pijuan-Sala et al., 2019). Each gene was manually categorized based on its enrichment in different tissues within this dataset. ‘No enrichment’ indicates genes that did not show a tissue-specific expression or enrichment. Arrow highlights *Afp* and arrowheads highlight genes associated with the complement and coagulation cascades. **C.** Confocal MIPs of immunostained embryos *Afp*-GFP; *Rreb1*^+/+^ and *Rreb1*^−/−^ embryos. Arrowheads mark highlight the proximal ExVE that, in contrast to wild-type embryos, shows little to no *Afp*-GFP expression. Sb, 50 μm. **D.** Brightfield images of E9.5 and 10.5 embryos showing abnormal defects in the vasculature of *Rreb1*^−/−^ embryos. In panel vi, arrowhead highlights the open anterior neural tube. **E.** Confocal maximum intensity projections of whole E9.5 and 10.5 embryos (Sb, 200 μm) with adjacent high magnification images of the cranial vasculature (Sb, 50 μm). Boxes i-iv in E9.5 are shown at higher magnification in Figure S3G. PECAM-1 marks vasculature. ENDOGLIN marks endothelial cells as well as hematopoietic, mesenchymal and neural stem cells. To note, the tail of the lower right embryo was damaged during dissection. **F.** Wholemount image of an E10.5 *Rreb1*^LacZ/LacZ^ mutant embryo. Arrowhead highlights pericardial edema. A, anterior; P, posterior; Pr, proximal; Ds, distal; D, dorsal; V, ventral; L, left; R, right; ExVE, extraembryonic VE; EmVE, embryonic VE; ys, yolk sac; fb, forebrain; mb, midbrain; hb, hindbrain; BVs, blood vessels; BCs, blood cells.

To assess the function of these genes, we implemented Gene Ontology (GO) and Kyoto Encyclopedia of Genes and Genomes (KEGG) pathway analysis. Downregulated genes were enriched for multiple GO terms associated with blood, including ‘blood microparticle’, ‘fibrinogen complex’, and ‘platelet alpha granule’ (Table S2), and the ‘complement and coagulation cascades’ (Table S3) that play a role in vasculogenesis (Girardi, Yarilin, Thurman, Holers, & Salmon, 2006; Moser & Patterson, 2003). Key genes within these groups included the complement inhibitor proteins *Cd59a*, and complement component factor I (*Cfi*), and the secreted proteins fibrinogen alpha and gamma (*Fga*, *Fgg*), complement factor B (*Cfb*), protein C (*Proc*) and *Alpha fetoprotein* (*Afp*). We also observed a downregulation of *Jag2* and *Slit1* (Table S1), components of the Notch and Slit-Robo signaling pathways respectively that regulate hematopoiesis and vasculogenesis (Blockus & Chedotal, 2016; Kofler et al., 2011).

The majority of these factors (84% of 55 transcripts detected by scRNA-seq of gastrulating mouse embryos (Nowotschin et al., 2019; Pijuan-Sala et al., 2019)) were specifically expressed by or highly enriched within the VE (Figure 3B, S3A). As our data was generated by whole embryo bulk RNA-sequencing, the downregulation, almost solely, of VE-associated genes could represent a relative decrease in the size of the VE compared to other tissues. However, other critical VE-associated genes, for example, the transcription factors and VE lineage determinants *Gata6*, *Gata4*, *Sox17*, and *Hnf4α* (Figure S3B), were not altered in *Rreb1*^−/−^ mutants. Thus, the observed transcriptional changes did not represent a global shift in the VE program.

*Afp* (downregulated in *Rreb1*^−/−^) is a plasma glycoprotein secreted by the yolk sac and fetal liver that regulates angiogenesis (O. D. Liang et al., 2004; Takahashi, Ohta, & Mai, 2004). To validate our RNA-sequencing, we crossed *Rreb1*^+/−^ mice to a transgenic reporter whereby the *Afp* cis-regulatory elements drive GFP expression (Kwon et al., 2006), and analyzed *Rreb1*^+/+^ and *Rreb1*^−/−^; *Afp*-GFP^*Tg*/+^ embryos (Figure S3C). In wild-type E7.5 and 8.5 embryos, *Afp*-GFP is expressed by embryonic VE cells (Kwon et al., 2006; Kwon, Viotti, & Hadjantonakis, 2008), and throughout the extraembryonic VE, with highest levels at the embryonic-extraembryonic boundary (Figure 3C, S3D). Like their wild-type littermates, *Rreb1*^−/−^ mutant embryos expressed *Afp*-GFP within the embryonic VE and at the embryonic-extraembryonic boundary but, consistent with our transcriptional data, showed little to no *Afp*-GFP within the extraembryonic VE (Figure 3C, S3D).

Based on these transcriptional changes, we then asked whether vascular development was perturbed in the absence of *Rreb1*. We observed that, at E9.5-10.5, wild-type embryos established a hierarchical branched network of blood vessels within the yolk sac and embryo-proper (Figure 3Di-iii) but, in contrast, *Rreb1*^−/−^ embryos had dysmorphic yolk sac capillaries that resembled a primitive capillary plexus (5/6 *Rreb1*^−/−^ at E9.5, 4/6 *Rreb1*^−/−^ at E10.5, Figure 3D iv, S3E, F i) and leakage of blood into the extravascular space (Figure 3D v, S3E). Various cardiovascular defects were also observed within the *Rreb1*^−/−^ embryo-proper including little to no blood within the fetus (2/6 *Rreb1*^−/−^ at E9.5, 1/6 *Rreb1*^−/−^ at E10.5, Figure 3D vi, S3F iii), pooling of blood (2/6 *Rreb1*^−/−^ at E9.5, 4/6 *Rreb1*^−/−^ at E10.5), a reduced vascular network (3/6 *Rreb1*^−/−^ at E10.5, Figure S3F ii) with fewer and wider blood vessels, particularly apparent in the cranial region (Figure 3E, S3G, H), widespread hemorrhaging (1/6 *Rreb1*^−/−^ at E9.5, Figure 3D vii), an enlarged heart (2/6 *Rreb1*^−/−^ at E9.5, Figure 3D vi), and pericardial edema (2/8 *Rreb1*^−/−^ at E10.5, Figure 3F). Therefore, loss of *Rreb1* results in the downregulation of vasculogenesis-associated genes and compromised cardiovascular development, culminating in embryonic lethality at midgestation.

### *Rreb1* regulates cytoskeleton and adherens junction organization within the epiblast

We then performed GO analysis of genes that were significantly upregulated in E7.5 *Rreb1*^−/−^ vs. *Rreb1*^+/+^ embryos, finding that these were enriched for 4 main categories; ‘cytoskeleton’, ‘membrane and vesicle trafficking’, ‘cell junctions’, and ‘extracellular space’ (Table S2). Factors associated with the cytoskeleton included microtubule components (*Tubb3*), microtubule-interacting proteins (*Map6*, *Jakmip2*, *Fsd1*), microtubule motors (*Kif5a*, *Kif5c*, *Kif12*), actin-binding proteins (*Coro1a*), and factors that connect adherens junctions to the cytoskeleton (*Ctnna2*, *Ablim3*) (Figure 4A). Genes within the ‘vesicle trafficking’ category were also related to the cytoskeleton. For example, Rab family members (*Rab6b*, *Rab39b*) (Figure 4A) regulate vesicle transport along actin and microtubule networks.

**Figure 4.**
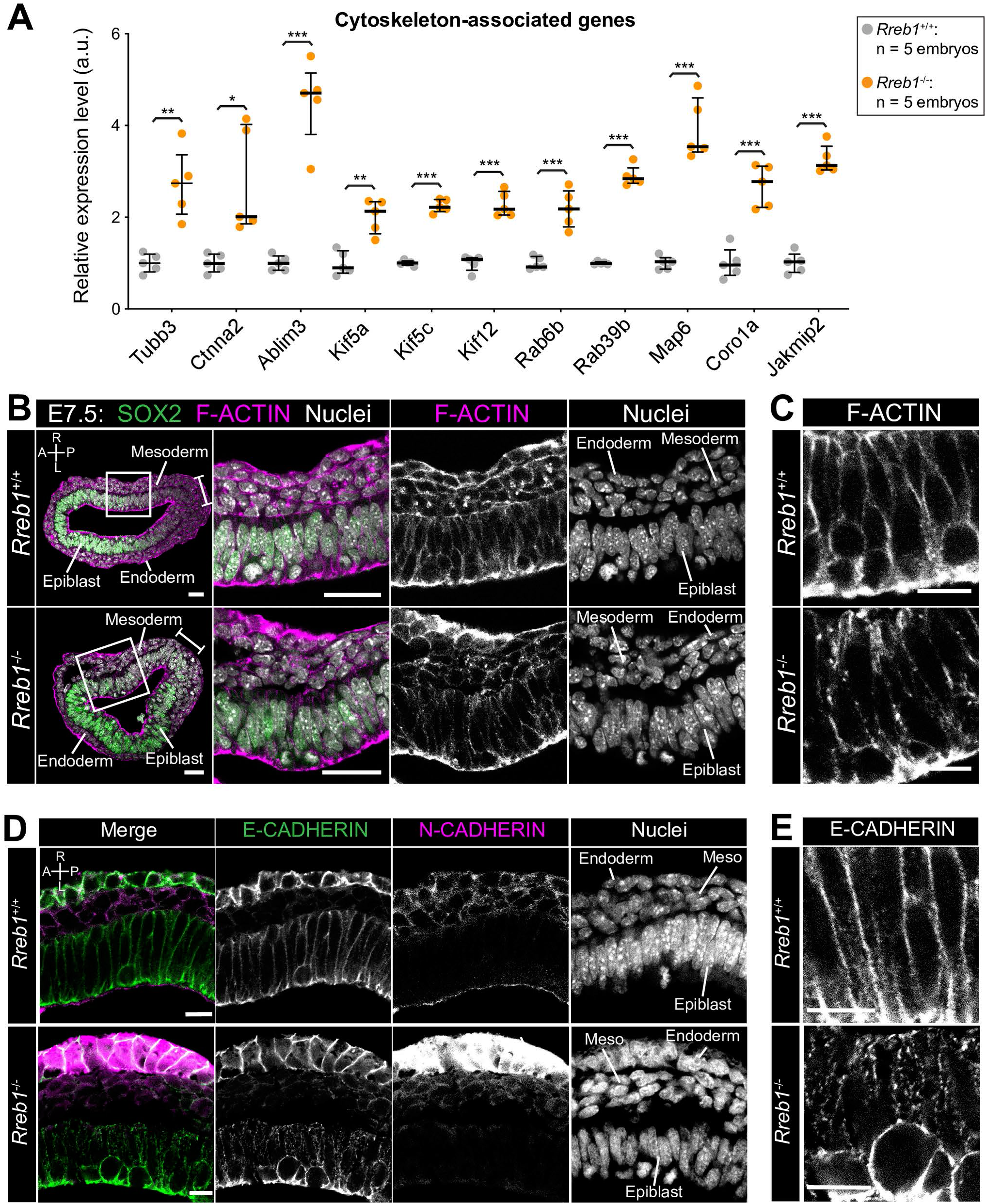
The *Rreb1*^−/−^ epiblast shows altered cytoskeleton and adherens junction organization. **A.** Graph showing the relative expression level of cytoskeleton-associated genes from RNA-sequencing of individual *Rreb1*^+/+^ and *Rreb1*^−/−^ embryos. Each point represents a single embryo. Statistical analysis was performed using an Unpaired *t*-test (\**p*<0.05, \*\**p*<0.005, \*\*\**p*<0.001). Bars represent median and IQR. Expression is shown relative to the mean expression in wild-type embryos. **B-E.** Confocal optical sections showing transverse cryosections of immunostained *Rreb1*^+/+^ and *Rreb1*^−/−^ embryos. Boxes indicate lateral epiblast regions shown at higher magnification in adjacent panels. *Rreb1*^−/−^ embryos exhibit a punctate localization of E-CADHERIN. Sb, 10 μm. **C,E.** Highest magnification images showing a small region of the epiblast epithelium. Sb, 10 μm. Brackets mark the primitive streak. A, anterior; P, posterior; L, left; R, right.

We therefore asked whether these cytoskeleton-centered transcriptional changes corresponded to a change in cytoskeleton organization in *Rreb1*^−/−^ mutants. In the normal (wild-type, *Rreb1*^+/+^) epiblast epithelium F-ACTIN was arranged into linear filaments oriented parallel to cell junctions (Figure 4B, C). In contrast, we found that F-ACTIN was punctate at proximal epiblast cell junctions within *Rreb1*^−/−^ embryos (Figure 4B, C).

The cytoskeleton interacts with and influences the localization of adherens junction components (X. Y. Chen, Kojima, Borisy, & Green, 2003; X. Liang, Gomez, & Yap, 2015; Mary et al., 2002; Mege & Ishiyama, 2017; Sako-Kubota, Tanaka, Nagae, Meng, & Takeichi, 2014; Stehbens et al., 2006; Teng et al., 2005). As we noted a significant upregulation of *Ctnna2* and *Ablim3*, which encode proteins connecting the cytoskeleton to adherens junctions (Figure 4A), we asked whether the change in F-ACTIN was also associated with a rearrangement of cell junctions. Cadherins are critical components of adherens junctions and, during gastrulation, E-CADHERIN is expressed within the epiblast, VE, and extraembryonic ectoderm (Pijuan-Sala et al., 2019). In wild-type embryos, E-CADHERIN, similar to F-ACTIN, forms a continuous belt between epithelial epiblast cells but, in *Rreb1* mutants, showed a punctate localization (Figure 4D, E, S4A-C). β-CATENIN was also more punctate at *Rreb1* mutant compared to wild-type epiblast junctions (Figure S4D), indicating that adherens junction complexes were altered. The change in E-CADHERIN and β-CATENIN protein localization was not associated with a transcriptional change in these genes, or in the expression of other adhesion-associated factors, such as tight junction components (Figure S4E), indicating that this altered localization occurs by post-transcriptional mechanisms. Therefore, loss of *Rreb1* results in a change in the expression of cytoskeleton-associated factors and a change in the organization of the cytoskeleton and adherens junctions within the epiblast.

### *Rreb1* maintains epithelial architecture of embryonic and extraembryonic tissues

The cytoskeleton is the scaffold of the cell that regulates cell-cell adhesion (Elson, 1988; Gavara & Chadwick, 2016; Grady, Composto, & Eckmann, 2016; Ketene, Roberts, Shea, Schmelz, & Agah, 2012) and epithelial organization (Bachir, Horwitz, Nelson, & Bianchini, 2017; Ivanov, Parkos, & Nusrat, 2010; B. Sun, Fang, Li, Chen, & Xiang, 2015; Vasileva & Citi, 2018). In cancer, a cytoskeleton-mediated switch from linear to punctate E-CADHERIN can occur, resulting in weaker cell-cell adhesion and loss of epithelial integrity (Aiello et al., 2018; Ayollo, Zhitnyak, Vasiliev, & Gloushankova, 2009; Gloushankova, Rubtsova, & Zhitnyak, 2017; Jolly et al., 2015; Kovac, Makela, & Vallenius, 2018; Saitoh, 2018). In keeping with this, *Rreb1*^−/−^ embryos exhibited perturbed epithelial architecture during gastrulation. In wild-type embryos, VE cells form an ordered monolayer epithelium overlying the embryonic epiblast and the ExE (Figure 5A, B, S5A), while in *Rreb1*^−/−^ embryos, cells protruded from the VE at various angles (Figure 5A), and the extraembryonic VE was frequently ruffled (Figure 5B, S5A). Moreover, abnormal masses of E-CADHERIN+ VE cells accumulated at the anterior embryonic-extraembryonic boundary (Figure 5C, S5B).

**Figure 5.**
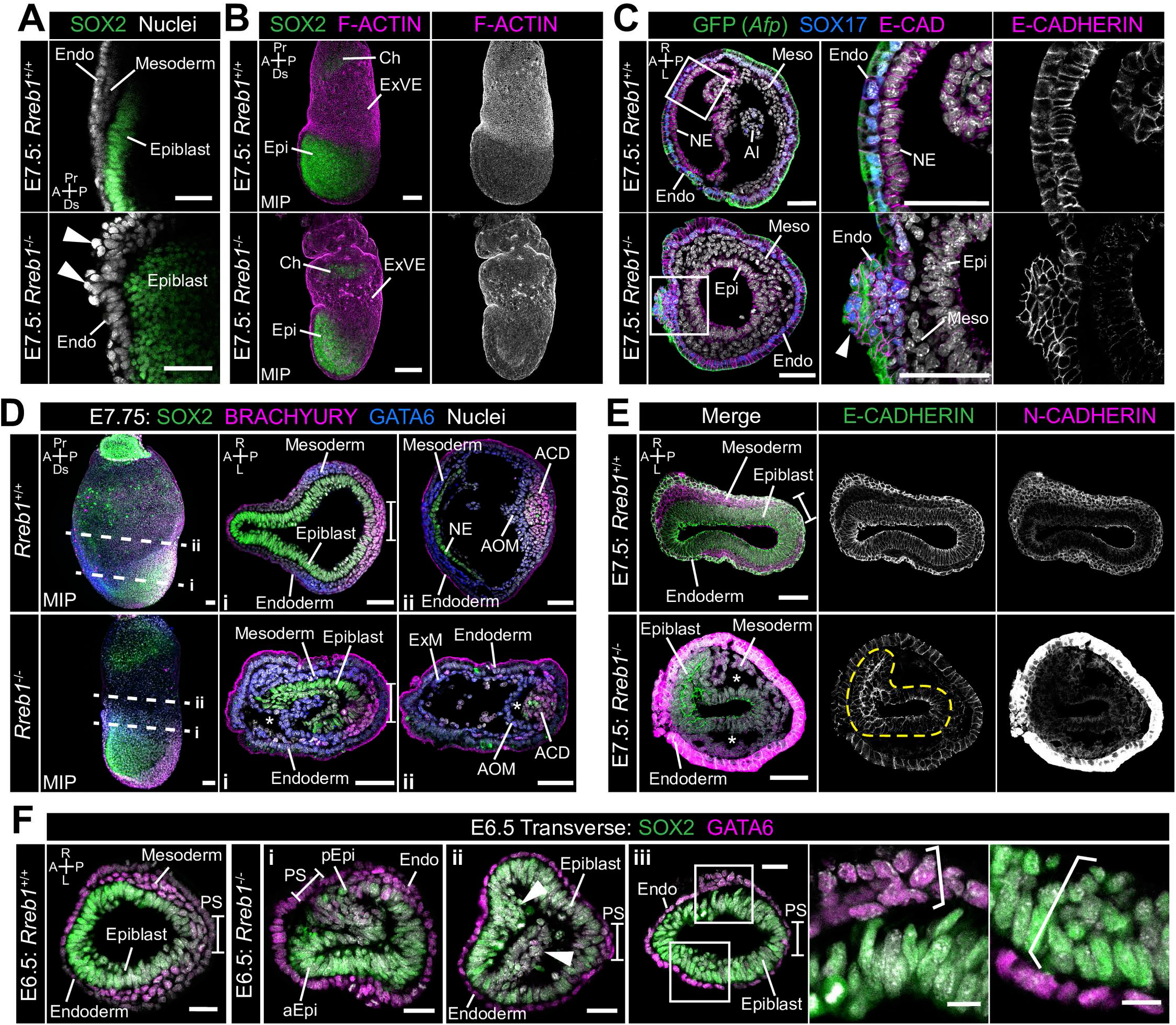
*Rreb1* maintains epithelial organization in the early mouse embryo. **A.** Sagittal confocal optical section of the anterior of E7.5 *Rreb1* wild-type and homozygous mutant embryos. Arrowheads highlight cells abnormally protruding from the VE overlying the epiblast. Sb, 25 μm. **B.** Confocal maximum intensity projections (MIP) of immunostained E7.5 embryos showing ruffling of the extraembryonic VE. Sb, 100 μm. **C.** Confocal optical sections showing transverse cryosections of E7.5 *Afp*-GFP *Rreb1* wild-type and homozygous mutant embryos. Boxes indicate regions shown in higher magnification in adjacent panels. Arrowhead indicates abnormal accumulation of *Afp*+ VE cells and underlying *Afp*-DE cells at the anterior embryonic-extraembryonic boundary in *Rreb1*^−/−^. Sb, 50 μm. **D,E.** Maximum intensity projections (MIPs) of wholemount E7.5 embryos and confocal optical sections of transverse cryosections. **D.** Dashed lines mark approximate plane of section. Sb, 50 μm. **E.** Dashed yellow line outlines the epiblast. Sb, 50 μm. Asterisks mark abnormal gaps between tissue layers. **F.** Representative images of *Rreb1*^+/+^ and *Rreb1*^−/−^ embryos highlighting the epithelial defects observed: (i) abnormal accumulations of cells in the epiblast, (ii) epiblast folding, in this case the epiblast is folded such that the putative anterior (aEpi) and posterior (pEpi) regions are adjacent to one another, (iii) formation of multilayered regions (highlighted with brackets) in the, typically monolayer, endoderm and epiblast. Sb 25 μm, high mag sb, 10 μm. **G-I.** Confocal MIPs (G,H) and confocal optical sections showing transverse cryosections of *Afp*-GFP; *Rreb1*^+/+^ and *Rreb1*^−/−^ embryos (I). Boxes indicate region shown in higher magnification in H. White circles indicate approximate embryonic-extraembryonic boundary. Sb, 50 μm. Pr, proximal; Ds, distal; A, anterior; P, posterior; R, right; L, left; Epi, epiblast; aEpi, anterior epiblast; pEpi, posterior epiblast; PS, primitive streak; Endo, endoderm; ACD, allantois core domain; AOM, allantois outer mesenchyme; Ch, chorion; Meso, mesoderm; ExVE, extraembryonic visceral endoderm; EmVE, embryonic visceral endoderm; DE, definitive endoderm; NE, neurectoderm; Al, allantois.

*Rreb1*^−/−^ initiated gastrulation in the posterior of the embryo, as marked by downregulation of the pluripotency-associated transcription factor SOX2 and upregulation of the primitive streak marker BRACHYURY (Figure 5D). Furthermore, *Rreb1*^−/−^ epiblast cells underwent an EMT at the primitive streak, delaminated from the epithelium, and migrated anteriorly in the wings of mesoderm (Figure 5E). While there was an increase in the fluorescence intensity of N-CADHERIN immunostaining within the VE of *Rreb1*^−/−^ vs. *Rreb1*^+/+^ embryos (Figure 5E), this was also observed with other antibodies and is likely associated with changes in the architecture of the VE leading to an increase in non-specific background staining within this tissue. Cells within *Rreb1*^−/−^ embryos also differentiated into mesoderm and DE, marked by GATA6 and SOX17 expression respectively (Figure 5D, S5C). Hence, *Rreb1*^−/−^ mutant cells can specify and begin to pattern the embryonic germ layers.

However, the mutant epiblast showed a range of morphological defects similar to those within the VE, including uncharacteristic folding of the epithelial layer (Figure 5E, F i, S5D), abnormal accumulations of cells (Figure 5F ii), increasingly variable cell orientation (Figure S5E-G), separation of typically closely apposed tissue layers, such as the mesoderm and endoderm (Figure 5D, E, S5H), and cells falling out of the epiblast (Figure S5I). In wild-type embryos, epiblast cells divide at the apical, cavity-facing surface while being maintained within the epithelial layer but, in *Rreb1*^−/−^ embryos, we observed dividing cells that left the epithelium (Figure S5J). Additionally, the epiblast and endoderm are monolayer epithelia in wild-type embryos but formed multilayered regions in *Rreb1*^−/−^ mutants (Figure 5F iii).

Epithelial homeostasis requires tight regulation of proliferation and the maintenance of cell polarity. *Rreb1*^−/−^ embryos showed no difference in the absolute or relative number of dividing cells within the epiblast, VE, or mesoderm when compared to wild-type littermates (Figure S5K, L). Furthermore, apicobasal polarity of the *Rreb1*^−/−^ epiblast cells was unaffected, demonstrated by the correct positioning of the tight junction protein ZO-1 at the apical surface and the basement membrane protein LAMININ at the basal surface (Figure S5M, N). Together these data show that loss of *Rreb1* results in disrupted epithelial architecture of both embryonic and extraembryonic tissues, associated with altered cytoskeleton and adherens junction organization.

### *Rreb1* mutant embryos display invasive phenotypes

In the context of cancer, cells that display punctate E-CADHERIN localization are considered to represent an intermediate epithelial-mesenchymal state (Sha et al., 2019; Yang et al., 2020), characterized by an increased propensity for collective invasion and metastasis (Aiello et al., 2018; Ayollo et al., 2009; Gloushankova et al., 2017; Jolly et al., 2015; Kovac et al., 2018; Saitoh, 2018). This state is linked to the downregulation of the transcription factor *Ovol1* (Jia et al., 2015; Saxena, Srikrishnan, Celia-Terrassa, & Jolly, 2020), which suppresses a mesenchymal identity, and the tight junction component, *Claudin7* (Aiello et al., 2018; W. K. Kim et al., 2019; Wang, Xu, Li, & Ding, 2018). Notably, both of these factors were also significantly downregulated in *Rreb1*^−/−^ embryos (Figure S6A). Furthermore, we observed that some *Rreb1*^−/−^ epiblast cells acquired mesenchymal characteristics. In wild-type embryos, the mesenchymal marker and EMT regulator SNAIL was expressed within the primitive streak and the wings of mesoderm (Figure 6A). However, in *Rreb1*^−/−^ embryos SNAIL was ectopically expressed within epiblast cells that were precociously exiting the epithelium (Figure 6A). Moreover, these cells exhibited punctate β-CATENIN, in contrast to the linear localization observed in neighboring SNAIL negative epiblast cells (Figure S6B). In *Rreb1*^−/−^ embryos, we also occasionally observed chains of cells that traversed tissue layers, including cells expressing the epiblast marker SOX2 that crossed the VE (Figure 6B, S6C) and cells expressing the mesoderm and endoderm marker GATA6 that spanned the epiblast (Figure S6C). In the majority of cases, these aberrant cells crossed the VE (Figure S6C, D) and SOX2-positive (SOX2+) pyknotic nuclei were detected on the adjacent exterior surface of the embryo (Figure S6C).

**Figure 6.**
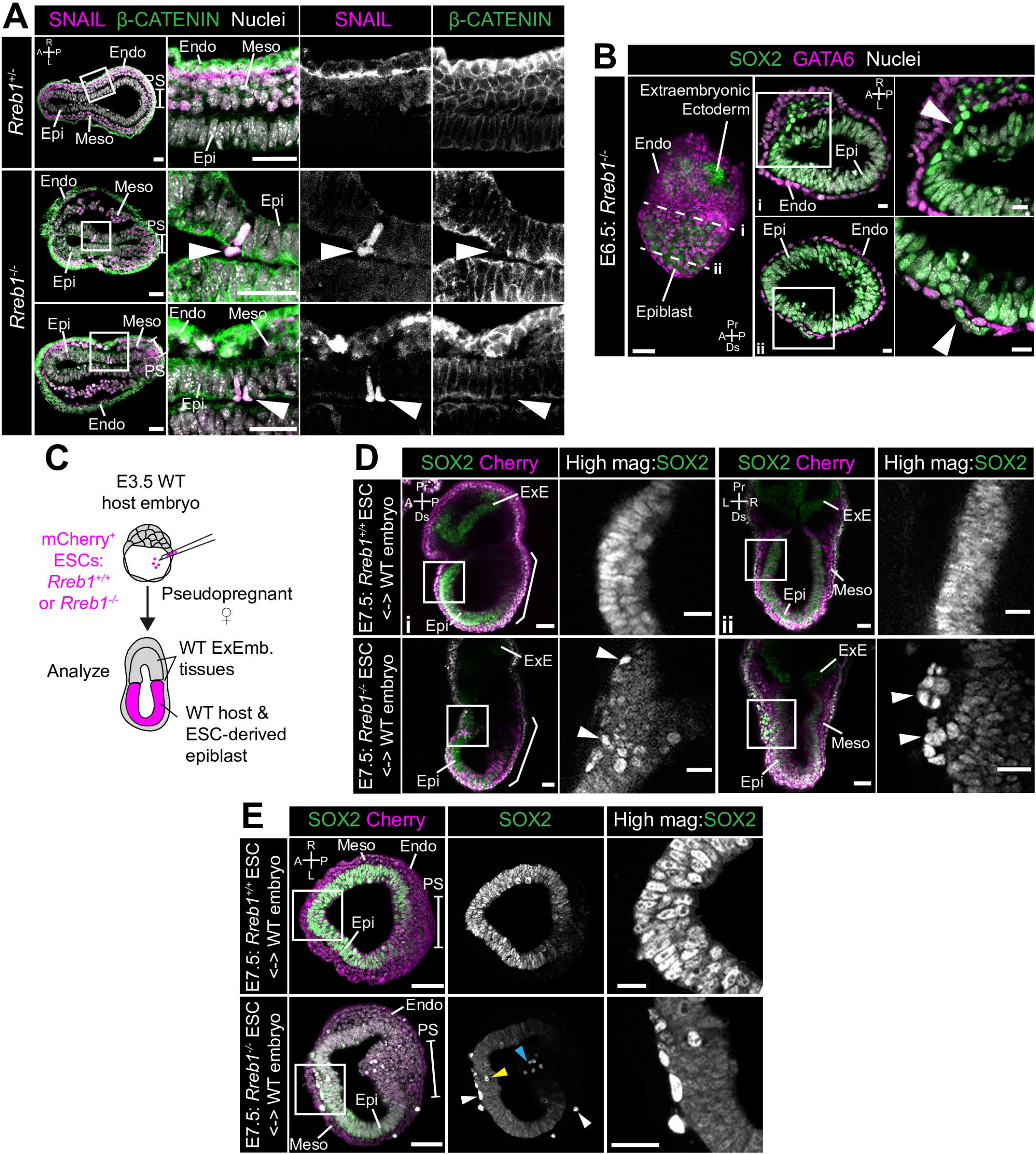
Loss of Rreb1 results in invasive cell behaviors. **A.** Confocal optical sections of transverse cryosections of immunostained E7.5 embryos. Boxes show regions displayed at higher magnification in adjacent panels. Arrowheads indicate ectopic SNAIL expression in epiblast cells exiting the epithelium. Sb, 25 μm. **B.** Confocal optical sections of maximum intensity projection (MIP, Sb, 50 μm) and transverse cryosections of immunostained E6.5 *Rreb1*^−/−^ embryos. Dashed lines mark approximate plane of transverse section. Arrowhead marks ectopic SOX2+ cells leaving the epiblast and traversing the outer endoderm layer. Sb, 10 μm. **C.** Schematic diagram illustrating how chimeras were generated. *Rreb1*^+/+^ or *Rreb1*^−/−^ embryonic stem cells (ESCs) constitutively expressing an mCherry lineage label were injected into wild host E3.5 embryos. Embryos were then transferred to pseudopregnant host females and dissected for analysis at later developmental stages. **D,E.** Sagittal (D i), lateral (D ii) and transverse (E) confocal optical sections of immunostained E7.5 chimeric embryos containing either *Rreb1*^+/+^ or *Rreb1*^−/−^ cells. Arrowheads mark abnormal SOX2+ cells, expressing higher levels of SOX2 than their neighbors, in the epiblast (yellow), primitive streak (blue arrowhead) or between the epiblast and visceral endoderm layers (white). Sb, 50 μm. High magnification inset Sb, 25 μm. A, anterior; P, posterior; L, left; R, right; Endo, endoderm; Meso, mesoderm; Epi, epiblast; PS, primitive streak.

In order to examine the cell-autonomous versus non-cell-autonomous effects of *Rreb1*, we then generated chimeric embryos by introducing *Rreb1*^−/−^ ESCs into wild-type host embryos so that the embryonic epiblast-derived tissues are a mosaic of wild-type and mutant origin and extraembryonic tissues are wild-type (Figure 6C). In E7.5 chimeric embryos, we frequently observed ectopic SOX2+ cells were dispersed throughout the embryo (30/63, 48% of *Rreb1*^−/−^ chimeric embryos, Figure 6D, E, S6E, 33-190 ectopic SOX2+ cells/per embryo). These cells expressed higher levels of SOX2 than most cells within the epiblast epithelium (Figure S6F) and were predominantly sandwiched between the epiblast and outer endoderm (Figure S6G).

SOX2-high cells were also found less frequently within the epiblast, cavity, and wings of mesoderm (Figure S6G). These SOX2+ cells divided and persisted until later stages of development (Figure S6H). Ectopic cells emerged prior to, or at the onset of, gastrulation (Figure S6I), and hence this was not a secondary consequence of gastrulation defects (Su et al., 2020). Thus, loss of *Rreb1* causes cells to ectopically exit the pluripotent epiblast epithelium in gastrulating mouse embryos.

### Invasive cells in *Rreb1*^−/−^ chimeras are associated with a distinct ECM organization

In chimeric embryos, ectopic SOX2+ cells were of both wild-type and mutant origin (Figure 7A, S7A), indicating that invasive-like behaviors were not driven solely by cell-autonomous properties, such as changes in the cytoskeleton and adherens junctions. Remodeling of the extracellular matrix (ECM) could promote invasive behaviors of both wild-type and mutant cells.

**Figure 7.**
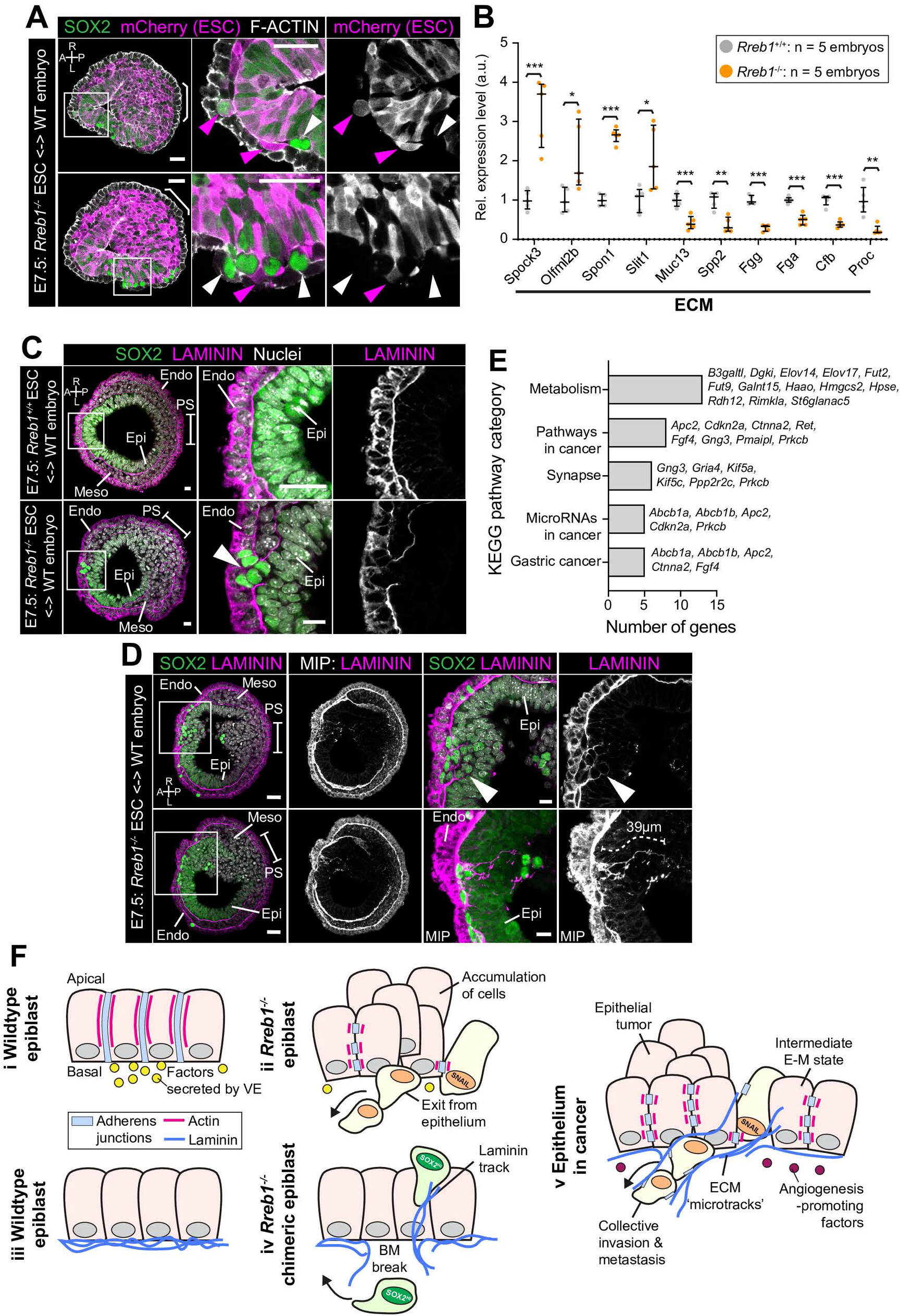
*Rreb1*^−/−^ chimeras exhibit changes in ECM organization. **A,C,D.** Confocal images showing transverse cryosections of immunostained E7.5 chimeric embryos containing *Rreb1*^+/+^ or *Rreb1*^−/−^ cells. **A.** Confocal optical sections of *Rreb1*^−/−^ chimeras. Cherry fluorescence is a constitutive lineage label marking the progeny of *Rreb1*^−/−^ embryonic stem cells (ESCs) introduced into host embryos. Arrowheads mark ectopic SOX2+ cells derived from wild-type host cells (white) or from *Rreb1*^−/−^ cells (magenta). Sb, 25 μm. **C.** Confocal optical sections of *Rreb1*^−/−^ chimeras. Arrowhead marks ectopic SOX2+ cells traversing a break in the basement membrane between the epiblast and outer visceral endoderm layer. Sb, 10 μm. **D.** Confocal optical sections and maximum intensity projections (MIP) of *Rreb1*^−/−^ chimeras. Upper and lower panels are sections taken from the same embryo, 20 μm apart. Arrowheads mark invasive SOX2+ cells surrounded by Laminin. Dashed line marks the approximate line of measurement of the length of the adjacent Laminin track. Sb, 25 μm and 10 μm for high magnification image. **B.** Graph showing the relative expression level of a panel of ECM-associated genes from RNA-sequencing of individual *Rreb1*^+/+^ and *Rreb1*^−/−^ embryos. Each point represents a single embryo. Statistical analysis was performed using an Unpaired *t*-test (\**p*<0.05, \*\**p*<0.005, \*\*\**p*<0.001). Bars represent median and IQR. Expression is shown relative to the mean expression in wild-type embryos. **E.** Graph showing the top 5 results from KEGG pathway analysis of genes that were significantly upregulated in *Rreb1*^−/−^ versus *Rreb1*^+/+^ embryos. The genes associated with each category are shown on the graph. **F.** Schematic diagram summarizing some of the key findings in this paper. i. In the wild-type epiblast epithelium of the mouse embryo, adherens junction components, such as E-CADHERIN, form continuous belts along cell junctions and F-ACTIN forms linear filaments that run parallel to these junctions. ii. In *Rreb1*^−/−^ embryos, there was a reduction in the expression of a cohort of factors secreted by the VE, which may alter the behavior of epiblast cells. Furthermore, we observed various phenotypes in the *Rreb1*^−/−^ epiblast epithelium including a more variable cell orientation compared to that of wild-type embryos, abnormal accumulations of cells, ectopic expression of the mesenchymal marker SNAIL, and chains of cells apparently exiting the epithelial layer. iii. The wild-type epiblast epithelium forms a Laminin basement membrane at its basal surface. iv. In contrast, in chimeric embryos that contain a mix of both wild-type and *Rreb1*^−/−^ cells, we observed cells of both genotypes traversing breaks in the underlying basement membrane which were then found ectopically throughout the embryo. Moreover, we observed the formation of long Laminin tracks closely associated with abnormal SOX2^HI^ cells. v. The cell behaviors observed in *Rreb1*^−/−^ embryos and chimeras are similar to those observed in cancer. For example, abnormal accumulations of epithelial cells are the basis of tumor formation, changes in cytoskeleton organization combined with a switch from linear to punctate E-CADHERIN and ectopic expression of mesenchymal markers characterizes an intermediate EMT state that is associated with collective invasion during cancer metastasis. Remodeling of the ECM into parallel fibers, known as ECM microtracks, facilitates collective cell invasion in cancer metastasis. Furthermore, the tumor microenvironment commonly show a change in the expression of secreted factors that promote angiogenesis. A, anterior; P, posterior; L, left; R, right; Pr, proximal; Ds, distal; Epi, epiblast; Endo, endoderm; ExE, extraembryonic ectoderm; Meso, mesoderm.

We noted that many of the genes that were significantly altered in *Rreb1*^−/−^ embryos were associated with ECM and cell-ECM adhesion. For example, *Tff3* (Ahmed, Griffiths, Tilby, Westley, & May, 2012; Pandey et al., 2014), *Hpse* (Liu et al., 2019), *Slit1 (Gara et al., 2015)*, *Spon1 (Chang et al., 2015)*, *Spock1* and *Spock3* (Q. Chen et al., 2016) are associated with increased cancer cell invasion and were upregulated in *Rreb1*^−/−^, and *Selenbp1* (Caswell et al., 2018; Schott et al., 2018) and *Serpin6b* (Chou et al., 2012) are tumor suppressor genes that were downregulated. including *Spock3*, *Spon1*, *Muc13*, and *Spp2* (Figure 7B, Table S2). Therefore, we asked whether the basement membrane underlying the epiblast was perturbed in *Rreb1*^−/−^ chimeras.

In wild-type chimeras, the basement membrane at the epiblast-VE interface is broken down in the posterior of the embryo at the primitive streak during gastrulation, as cells undergo an EMT (Figure 7C). In *Rreb1*^−/−^ embryo chimeras, the basement membrane was broken down at the primitive streak but also in anterior and lateral regions of the epiblast (Figure 7C, S7B). SOX2+ cells were observed traversing these ectopic basement membrane breaks (Figure 7C). Furthermore, aberrant SOX2+ cells were surrounded by higher levels of Laminin than their neighbors and associated with Laminin tracks, up to 68 μm (approximately 7 cell diameters) in length (Figure 7D, S7C). Thus, loss of *Rreb1* in the mouse embryo caused epiblast epithelial cells to cross the basement membrane underlying the epiblast epithelium, reminiscent of the invasive cell behaviors observed in cancer metastasis. These defects were associated with cell-autonomous changes in the cytoskeleton as well as non-cell-autonomous changes in the ECM. KEGG pathway analysis also revealed that the genes upregulated in *Rreb1*^−/−^ embryos were enriched for pathways associated with cancer, including ‘Pathways in cancer’, ‘MicroRNAs in cancer’, and ‘Gastric cancer’ (3/5 most enriched pathways, Figure 7E). Together these data suggest that the embryonic role of *Rreb1* may be functionally linked to its role in cancer (Figure 7F).

## 3. Discussion

The transcription factor *Rreb1* is necessary for invertebrate development (Melani et al., 2008; Pickup et al., 2002; Wieschaus et al., 1984; Wilk et al., 2000) and is implicated in cancer (Ferrara & De Vanna, 2016; Hui et al., 2019; Kent et al., 2017; Li et al., 2018; Mukhopadhyay et al., 2007; Thiagalingam et al., 1996; Yao et al., 2019), suggesting that it plays critical contextual organismal functions. Despite this, we know little about its role in mammalian development.

Here we demonstrate that *Rreb1* is essential for mouse embryo development. Loss of *Rreb1* resulted in disrupted epithelial architecture of both embryonic and extraembryonic tissues. These defects were consistent with the reported role of the Drosophila homolog of Rreb1, Hindsight (hnt), which regulates cell adhesion during invertebrate development (Melani et al., 2008; Pickup et al., 2002; Wilk et al., 2000). Pluripotent epiblast cells fell out of their epithelial layer into the space between the epiblast and VE in both *Rreb1*^−/−^ mutant embryos and chimeras. Similarly, loss of hnt in the Drosophila retina causes cells to fall out of the epithelium into the underlying tissue layer (Pickup et al., 2002). Thus, Rreb1 is an evolutionarily conserved regulator of tissue architecture.

*Rreb1* homozygous mutant embryos die at midgestation due to a range of cardiovascular defects, including perturbed yolk sac vasculogenesis. Although *Rreb1* was not highly expressed by the yolk sac mesoderm, which will give rise to endothelial cells, it was robustly expressed by the overlying yolk sac endoderm (Figure S1G). The yolk sac endoderm is known to secrete factors that regulate cardiogenesis, vasculogenesis and hematopoiesis (Arai, Yamamoto, & Toyama, 1997; Belaoussoff, Farrington, & Baron, 1998; Byrd et al., 2002; Damert, Miquerol, Gertsenstein, Risau, & Nagy, 2002; Dyer, Farrington, Mohn, Munday, & Baron, 2001; Goldie, Nix, & Hirschi, 2008; Miura & Wilt, 1969; Wilt, 1965). Moreover, *Rreb1* mutants showed a significant downregulation of a cohort genes encoding secreted vasculogenesis-associated factors, as well as genes involved in vesicular transport that form part of the secretory pathway. Thus, the role of *Rreb1* in embryonic vasculogenesis is likely mediated via paracrine interactions with the VE.

We previously showed that, in a cancer model, *Rreb1* directly binds to the regulatory region of *Snai1* in cooperation with TGF-β activated SMAD transcription factors to induce the expression of SNAIL, which drives EMT (Su et al., 2020). Furthermore, mouse embryos containing *Rreb1*^−/−^ cells exhibit an accumulation of cells at the primitive streak, consistent with a disrupted gastrulation EMT (Su et al., 2020). These data suggested that *Rreb1* may be required for EMT in both development and disease contexts. However, upon closer examination we found that loss of *Rreb1* also disrupts epithelial architecture. We found that, in the mouse embryo, *Rreb1* is expressed not only in mesenchymal tissues, such as the primitive streak and mesoderm, but also within epithelial tissues such as the trophectoderm, VE and the notochord. Thus, Rreb1 does not drive EMT in all contexts. Likewise, in Drosophila, hnt exhibits context-dependent adhesion regulation. For example, loss of hnt in the trachea and retina disrupts epithelial architecture (Pickup et al., 2002; Wilk et al., 2000), while loss of hnt from border cells results in increased cell-cell adhesion (Melani et al., 2008). Thus, its function likely depends on the combination of factors and signaling activities present within any given cell where it is expressed.

Global transcriptional analysis of *Rreb1*^−/−^ embryos revealed that loss of *Rreb1* significantly alters the transcription of cytoskeleton-associated genes, including actin-binding proteins, microtubule components and microtubule motor proteins. Hnt also genetically interacts with and transcriptionally regulates cytoskeleton-associated genes, such as *chickadee* (*Profilin1*), which governs actin polymerization and depolymerization, the F-ACTIN crosslinker *karst* (*Alpha-actinin-1*), Actin-binding protein *jitterbug* (*Filamin A*), a microtubule motor *dynamitin* (*Dynactin2*) and *Rho1*, a GTPase that regulates cytoskeleton organization (Oliva et al., 2015; Wilk, Pickup, Hamilton, Reed, & Lipshitz, 2004). While the specific factors downstream of *Rreb1* and hnt are distinct, these data suggest a conserved role in cytoskeleton regulation. The transcriptional changes in cytoskeleton regulators corresponded to a change in the organization of the cytoskeleton and adherens junctions whereby wildtype epiblast cell junctions displayed a continuous, linear arrangement of F-ACTIN, E-CADHERIN and β-CATENIN, while *Rreb1*^−/−^ exhibited a punctate localization. ACTIN interacts with cadherins (M. K. L. Han & de Rooij, 2017) and thus may directly influence their localization. The cytoskeleton mediates vesicular trafficking, which can also regulate E-CADHERIN localization (Aiello et al., 2018; X. Y. Chen et al., 2003; Chung et al., 2014; X. Liang et al., 2015; Mary et al., 2002; Pilot, Philippe, Lemmers, & Lecuit, 2006; Sako-Kubota et al., 2014; Stehbens et al., 2006; Teng et al., 2005; Vasileva & Citi, 2018), and a large number of trafficking genes were upregulated in *Rreb1*^−/−^ embryos. Therefore, a combination of altered vesicle trafficking and/or direct changes in the cytoskeleton may regulate E-CADHERIN localization. As *Rreb1* is not expressed highly throughout the epiblast, these phenotypes are either due to a loss of low-level epiblast expression or mediated through paracrine interactions with the VE. Future tissue-specific ablations of *Rreb1*, and chromatin immunoprecipitation (ChIP) studies to identify direct targets, will distinguish between these possibilities.

A reduction in ACTIN stress fibers enhances the motility and deformability of cells and is associated with an invasive phenotype in cancer (Grady et al., 2016; Y. L. Han et al., 2020; Katsantonis et al., 1994; Suresh, 2007; Xu et al., 2012). Moreover, altered ACTIN organization (Gloushankova et al., 2017; Kovac et al., 2018) and punctate E-CADHERIN is indicative of an intermediate epithelial-mesenchymal state, which also correlates with weaker cell-cell adhesion and collective invasion in metastasis (Aiello et al., 2018; George, Jolly, Xu, Somarelli, & Levine, 2017; Jolly et al., 2015; Saitoh, 2018). In keeping with this, *Rreb1*^−/−^ cells displayed invasive phenotypes *in vivo* resulting in ectopic SOX2+ epiblast-like cells positioned throughout chimeric embryos. However, ectopic cells were of wild-type and mutant origin indicating that not only cell-autonomous properties, such as cytoskeletal organization, but also cell non-autonomous mechanisms drive this behavior. *Rreb1*/hnt phenotypically interacts with and transcriptionally regulates ECM-associated factors such as *viking* (*Col4a1*), *Cg25c* (*Col4a2*), *Mmp2* and *Adamts5* (Deady, Li, & Sun, 2017; Wang et al., 2017; Wilk et al., 2004). We also observed a change in the expression of ECM-associated factors in *Rreb1*^−/−^ embryos, some of which have been linked to changes in the metastatic potential of cells. Furthermore, KEGG pathway analysis of downregulated genes revealed that these were associated with the complement and coagulation cascades, which control a variety of processes, including ECM remodeling, and the corruption of this pathway is linked to cancer metastasis (Ajona, Ortiz-Espinosa, Pio, & Lecanda, 2019). Thus, changes in ECM composition in *Rreb1*^−/−^ embryos may drive invasive behaviors. Ectopic SOX2+ cells were associated with abnormal breaks in the basement membrane, elevated levels of Laminin, and Laminin tracks. These ECM tracks are reminiscent of bundles of parallel Collagen fibers, referred to as “microtracks”, observed in cancer. Microtracks are generated through ECM remodeling by invasive leader cells, which subsequently facilitates the migration of less invasive cells within the tumor (Gaggioli, 2008; Gaggioli et al., 2007; Poltavets, Kochetkova, Pitson, & Samuel, 2018). Intriguingly, ectopic SOX2+ cells of wild-type origin were adjacent to *Rreb1*^−/−^ cells. Thus, *Rreb1*^−/−^ cells might perform a role comparable to leader cells in cancer metastasis, remodeling the ECM to permit migration of wild-type neighbors.

In sum, we have identified cell behaviors and phenotypes in *Rreb1* mutant mouse embryos, which are reminiscent of those observed during cancer cell invasion, including loss of epithelial architecture, aberrant basement membrane breakdown, ECM remodeling, and ectopic exit of cells from an epithelium. The early mouse embryo is an experimentally tractable *in vivo* system to interrogate these phenotypes and thus, future studies of the function of *Rreb1* in development may also shed light on its role in metastasis and other diseases involving loss of epithelial integrity.

## 4. Materials and methods

### Generation and maintenance of mouse lines

Mice were housed under a 12 hr light-dark cycle in a specific pathogen-free room in the designated facilities of MSKCC. Natural matings were set up in the evening and mice were checked for copulation plugs the following morning. The date of vaginal plug was considered as E0.5. Genotyping was carried out at the time of weaning. Mice were outbred to CD1 animals and maintained on a mixed bred CD-1/129 Sv/C57BL6/C2J background in accordance with the guidelines of the Memorial Sloan Kettering Cancer Center (MSKCC) Institutional Animal Care and Use Committee (IACUC).

To generate the *Rreb1*^LacZ^ reporter mouse line, *in vitro* fertilization was performed using C57BL/6N-A^tm1Brd^ Rreb1^tm1a(EUCOMM)Wtsi^/WtsiPh (RRID:IMSR_EM:10996) sperm obtained from the European Conditional Mouse Mutagenesis Program (EUCOMM). The Tm1a (knockout-first) allele was genotyped by PCR using the following primers: *Rreb1* 5’ arm: CTTCTGTCCCAGAAGCTACATTGC, *Rreb1* 3’ arm: GGACAACGGTCACTGAGAAGATGG, Lar3: CAACGGGTTCTTCTGTTAGTCC and the protocol: Step1 - 95 °C for 3 min, Step 2–35x: 95 °C for 30 s, 63 °C for 30 s, 72 °C for 30 s, Step 3–72 °C for 3 min. This results in a wild-type allele amplicon band of 751 bp and a transgenic allele amplicon of 502 bp. Tm1a mice were then crossed with a Flp recombinase mouse line (Rodriguez et al., 2000) to remove the neomycin cassette and Exon 6, producing the Tm1b LacZ tagged null allele. *Rreb1*^LacZ/+^ embryos were analyzed by X-gal staining to determine the *Rreb1* expression pattern.

*Rreb1*^−/−^ mutant mice were generated by CRISPR-mediated genetic knockout. The CRISPR gRNAs used for deleting exon 6 of the *Rreb1* gene were designed using the approach of Romanienko et. al (Romanienko et al., 2016). The sequences of the guides are: crRNA#1: TATTATGAACTCCTCTGGAC, crRNA#2: AGTGTCTTCGAAAGAGCCAA, crRNA#3: CGTTACAACAAAGCACCCTT, crRNA#4: AGGAAAACTCGTAGTGGCAC. To initiate cleavage and subsequent deletion of the target locus in mice, guides were injected in pairs, either #1 and #3 or #2 and #4, into the pronuclei of mouse zygotes at a concentration of 50 ng/μl each, with 100 ng/μl purified Cas9 protein (PNABio, Newbury Park, CA), using conventional techniques (Behringer, Gertsenstein, Vintersten Nagy, & Nagy, 2014). Founder mice were analyzed for the deletion by PCR using the primers RREB2: GACACCTAGTCACCGAGGAAAC and RREB6: CTGTGGCAGATCTGGTAGGC. This primer pair is located outside of the gRNA cleavage sites, thereby revealing the size of the deletion based on the nucleotide length of the amplicon obtained. The wild type amplicon size is 1019bp. The deletion amplicons, if there had been a simple cut and rejoining, would be: Cr#1 and #3: 275bp. Cr#2 and #4: 456bp. Genotyping of the *Rreb1* locus was performed by PCR with primers RREB1_1: GTGACAGAGGGAACAGTGGG, RREB1_2: GACACCTAGTCACCGAGGAAAC, RREB1_3: GTGTCTGTGTTGTGCTGCA using the following protocol: Step1 - 94 °C for 3 min, Step 2–35x: 95 °C for 30 s, 64 °C for 90 s, 72 °C for 1 min, Step 3–72 °C for 5 min, resulting in a 358 bp amplicon for the wild-type allele and a 275 bp amplicon for the mutant allele. *Rreb1*^−/−^ mice were embryonic lethal at midgestation but no peri-natal lethality was observed for *Rreb1*^−/+^ mice. Therefore, the *Rreb1* mouse line was maintained and *Rreb1*^−/−^ embryos were obtained through heterozygous *Rreb1*^−/+^ intercrosses.

### Generation of chimeric embryos

Approximately 10-15 *Rreb1*^−/−^ ESCs, described in (Su et al., 2020), harboring a constitutive mCherry fluorescent lineage tracer were injected into E3.5 blastocysts (C57BL/6J, Jackson Laboratory, Bar Harbor, ME) as previously described (Su et al., 2020). Injected blastocysts were cultured in KSOM/AA (Millipore, Billerica, MA) at 37ºC in an atmosphere of 5% CO2 to allow for recovery of blastocyst morphology and then implanted into the uterine horns (up to ten embryos per horn) of E2.5 pseudopregnant females (C57BL/6J;CBA F1, Jackson Laboratory) using standard protocols. Chimeric embryos were recovered between E7.5-E9.5.

### Wholemount in situ hybridization

To produce the *Rreb1* riboprobes, RNA was isolated from pooled E12.5 CD1 mouse embryos using an RNeasy^®^ Plus Mini Kit (Qiagen, Hilden, Germany) and then used to generate cDNA with a QuantiTect Reverse Transcription Kit (Qiagen), as per manufacturer’s instructions. Primers (5’ UTR L: GGGCCTTTGTCTCATGCTCC, 5’ UTR R: CGCAGAATGTTTTCCTCAACAG) were designed against a unique 502 bp region within the *Rreb1* 5’ UTR and used to PCR amplify this fragment from E12.5 embryo cDNA. The PCR product was purified using a QIAquick^®^ PCR Purification Kit (Qiagen) and a TOPO™TA Cloning™ Kit (K461020, Thermo Fisher Scientific) used to introduce the fragment into a pCR™II-TOPO™ Vector and transformed into E.coli. Colonies were picked, expanded and the plasmid isolated for sequencing. A plasmid containing the correct sequence (5’-CGCAGAATGTTTTCCTCAACAGTTGACAATTTTAGGATAAATAGAACTTTAGAAAAATTACTA CTATCAATCATCTAAGTATTCCGAATAGGAAAAAAAGTCAAAATAAGTAAGGGACGCTGGAGCTACCTCAGTGAAGGGGAAAAAATATCCAATCCCACTTTTCTGTATTACATGTGTGGTAGCTAAAGAACTCCATAGAATGTTCAAAAAAAAAAAAAAAAGACGGCACTGAAGATTATCATGTCAAAGCACCAAGCTCATTACATCACTGTTACCTTAATGCAAAGTCCCACTTCTCCGGAATGGCCTCCATACTTAGAAACTCTTGGAACTTGTCAGGCAAAGGTTATGGGGAGGGAAGTGAAGGAGCCTATGACCACTGTCACTGTGTCTGATACATTTATTTACAGATAAGCCTTGGTGGCTCAGACCACAGGCACAGATTATATGGAAAGTAACAGCCTGTGACTTCTGAGACAAAGAATGGAGCATGAGACAA-3’) was selected, linearized and the dual promoter system within the pCR™II-TOPO™ Vector used to amplify and DIG label both a control sense and an antisense probe. Wholemount mRNA *in situ* hybridization was then carried out as previously reported (Conlon & Rossant, 1992).

### X-gal staining

X-gal staining of cells and embryos containing the *Rreb1*-LacZ reporter was performed using a β-Gal Staining Kit (K146501, Invitrogen, Waltham, MA) as per manufacturer’s instructions. Embryos and cells were fixed for 15 mins at room temperature followed by staining until the blue color was detectable (2-3 hours) at 37 °C.

### Cell culture

Cells were maintained in standard serum/LIF ESC medium (Dulbecco’s modified Eagle’s medium (DMEM) (Gibco, Gaithersburg, MD) containing 0.1 mM non-essential amino-acids (NEAA), 2 mM glutamine and 1 mM sodium pyruvate, 100 U/ml Penicillin, 100 μg/ml Streptomycin (all from Life Technologies, Carlsbad, CA), 0.1 mM 2-mercaptoethanol (Sigma, St. Louis, MO), and 10% Fetal Calf Serum (FCS, F2442, Sigma) and 1000 U/ml LIF) as previously described (Morgani, Metzger, Nichols, Siggia, & Hadjantonakis, 2018). C57BL/6N-A^tm1Brd^ Rreb1^tm1a(EUCOMM)Wtsi^/WtsiPh (RRID:IMSR_EM:10996) embryonic stem cell lines were used to analyze *Rreb1* expression and also converted to an epiblast stem cell (EpiSC) state through prolonged culture (more than 5 passages) in N2B27 medium containing 12 ng/ml FGF2 (233-FB-025, R&D Systems) and 20 ng/ml ACTIVIN A (120-14P, Peprotech, Rocky Hills, NJ), as previously described (Tesar et al., 2007).

### Immunostaining

Cell lines were immunostained as previously described (Morgani, Metzger, et al., 2018). Post-implantation embryos were fixed in 4 % paraformaldehyde (PFA) for 15 min at room temperature (RT). Embryos were washed in phosphate-buffered saline (PBS) plus 0.1 % Triton-X (PBST-T) followed by 30 min permeabilization in PBS with 0.5 % Triton-X. Embryos were washed in PBS-T and then blocked overnight at 4 °C in PBS-T, 1 % bovine serum albumin (BSA, Sigma) and 5 % donkey serum. The following day, embryos were transferred to the primary antibody solution (PBS-T with appropriate concentration of antibody) and incubated overnight at 4 °C. The next day, embryos were washed 3 x 10 min in PBS-T and then transferred to blocking solution at RT for a minimum of 5 hr. Embryos were transferred to secondary antibody solution (PBS-T with 1:500 dilution of appropriate secondary conjugated antibody) and incubated overnight at 4 °C. Embryos were then washed 3 x 10 min in PBS-T with the final wash containing 5 μg/ml Hoechst. Where F-ACTIN staining was performed, Alexa Fluor^TM^ conjugated phalloidin (Thermo Fisher Scientific, Waltham, MA) was added to the primary and secondary antibody solutions at a 1:500 dilution.

### Antibodies

The following primary antibodies were used in this study: β-catenin (RRID:AB_397555, BD Transduction labs, Billerica, MA, 610154, 1:500), Brachyury (RRID:AB_2200235, R&D, AF2085, 1:100), CD31 (RRID:AB_394819, BD Biosciences, 553373, 1:100) CD105 (RRID:AB_354735, R&D Systems, AF1320, 1:100), E-cadherin (RRID:AB_477600, Millipore Sigma, U3254, 1:200), Gata6 (RRID:AB_10705521, D61E4 XP, Cell Signaling, 5851, 1:500), GFP (RRID:AB_300798, Abcam, ab13970), Laminin (RRID:AB_477163, Millipore Sigma, L9393, 1:500), N-cadherin (RRID:AB_2077527, BD Biosciences, 610920, 1:200), RFP (Rockland, Limerick, PA, 600-400-379, 1:300), Snail (RRID:AB_2191738, R&D Systems, AF3639, 1:50), Sox2 (RRID:AB_11219471, Thermo Fisher Scientific, 14-9811-82, 1:200), Sox17 (RRID:AB_355060, R&D Systems, AF1924, 1:100), ZO-1 (RRID:AB_87181, Invitrogen, 33-9100, 1:200).

### Cryosectioning

Embryos were oriented as desired and embedded in Tissue-Tek^®^ OCT (Sakura Finetek, Japan). Samples were frozen on dry ice for approximately 30 min and subsequently maintained for short periods at −80 °C followed by cryosectioning using a Leica CM3050S cryostat.

Cryosections of 10 μm thickness were cut using a Leica CM3050S cryostat and mounted on Colorfrost Plus^®^ microscope slides (Fisher Scientific) using Fluoromount G (RRID:SCR_015961, Southern Biotech, Birmingham, AL) and imaged using a confocal microscope as described.

### Confocal imaging and quantitative image analysis

Embryos were imaged on a Zeiss LSM880 laser scanning confocal microscope. Whole-mount embryos were imaged in glass-bottom dishes (MatTek, Ashland, MA) in PBS. Raw data were processed in ImageJ open-source image processing software (Version: 2.0.0-rc-49/1.51d).

Nuclei orientation (Figure S5E-G) was measured manually using Fiji (RRID:SCR_002285, Image J) software. Using the angle tool, we measured the angle between the long axis of individual epiblast nuclei and the underlying basement membrane, marked by Laminin staining on confocal optical sections of transverse cryosections. We measured the angle of 143 cells from 3 *Rreb1*^+/+^ embryos and 136 cells from 3 *Rreb1*^−/−^ embryos.

We quantified proliferation in *Rreb1*^+/+^ versus *Rreb1*^−/−^ embryos (Figure S5L) by manually counting the number of phosphorylated histone H3 (pHH3) positive cells in the epiblast, outer endoderm layer or wings of mesoderm in transverse cryosections of *Rreb1*^+/+^or *Rreb1*^−/−^ embryos. Initially, cell counts were also categorized as divisions in anterior versus posterior embryonic regions but, as no differences were observed, these data were subsequently combined. We performed counts on cryosections comprising 3 entire embryos per genotype. Data was analyzed as the absolute numbers of dividing cells per cell type. Additionally, we counted the total number of cells per cell type per section and normalized the number of dividing cells to this value to account for differences based on embryo or tissue size. Statistics were performed on a per embryo rather than a per cell basis.

The level of GFP in the VE of *Afp*-GFP; *Rreb1*^+/+^ and *Rreb1*^−/−^ embryos was quantified by manually selecting the embryonic and extraembryonic region of confocal maximum intensity projection images and measuring the mean fluorescence intensity using Fiji software.

Quantification of SOX2 protein levels (Figure S6F) were carried out on cryosections of *Rreb1*^−/−^ chimeric embryos containing cells expressing high levels of SOX2 (SOX2^HI^ cells) to determine the approximate fold change in protein level relative to normal surrounding cells. To make measurements, nuclei were manually identified using the freehand selection tool in Fiji software. Aberrant SOX2^HI^ cells could readily be distinguished from standard neighboring cells by their elevated signal after immunostaining for SOX2 protein. Mean fluorescence intensity of SOX2 immunostaining was measured within all SOX2^HI^ nuclei within a particular cryosection and an equivalent number of randomly selected nuclei with normal SOX2 expression within the anterior and posterior epiblast regions were measured. Mean SOX2 fluorescence intensity in each nucleus was normalized to the corresponding mean fluorescence intensity of the Hoechst nuclear stain. All data is shown relative to the mean SOX2 fluorescence intensity measured in ‘normal’ anterior epiblast cells of the same confocal optical section. A total of 8 embryos, 35 cryosections and 696 cells were analyzed. Statistics were carried out on the average fluorescence levels per embryo.

The localization of SOX2^HI^ cells (identified manually from SOX2 immunostaining) (Figure S6G) was scored based on their location within confocal images of cryosectioned *Rreb1^−/−^* chimeric embryos. Scoring was carried out on 76 cryosections from 7 independent embryos that contained high numbers of SOX2^HI^ cells. SOX2^HI^ cells were scored as being within the Epi itself, at the Epi-VE interface (outside of the epiblast epithelium), within the primitive streak or wings of mesoderm (mesoderm) or within the amniotic cavity.

### Statistics

Statistical analysis of significance was assessed using a One-way ANOVA (p<0.0001) followed by unpaired *t*-tests to compare particular groups (GraphPad Prism, RRID:SCR_002798, GraphPad Software, Inc., Version 7.0a).

### RNA-sequencing and data analysis

Frozen tissue was homogenized in TRIzol Reagent (ThermoFisher catalog # 15596018) using the QIAGEN TissueLyser at 15 Hz for 2-3 min with a Stainless-Steel Bead (QIAGEN catalog # 69989). Phase separation was induced with chloroform. RNA was precipitated with isopropanol and linear acrylamide and washed with 75% ethanol. The samples were resuspended in RNase-free water. After RiboGreen quantification and quality control by Agilent BioAnalyzer, 150 g of total RNA underwent polyA selection and TruSeq library preparation according to instructions provided by Illumina (TruSeq Stranded mRNA LT Kit, catalog # RS-122-2102), with 8 cycles of PCR. Samples were barcoded and run on a HiSeq 4000 in a 50 bp/50 bp paired**-** end run, using the HiSeq 3000/4000 SBS Kit (Illumina). An average of 47 million paired reads was generated per sample. The percent of mRNA bases averaged 67%.

The output data (FASTQ files) were mapped to the target genome using the rnaStar aligner (Dobin et al., 2013) that maps reads genomically and resolves reads across splice junctions. We used the 2 pass mapping method outlined in (Engstrom et al., 2013), in which the reads are mapped twice. The first mapping pass uses a list of known annotated junctions from Ensemble. Novel junctions found in the first pass were then added to the known junctions and a second mapping pass is done (on the second pass the RemoveNoncanoncial flag is used). After mapping we post-processed the output SAM files using the PICARD tools to: add read groups, AddOrReplaceReadGroups which in additional sorts the file and converts it to the compressed BAM format. We then computed the expression count matrix from the mapped reads using HTSeq (www-huber.embl.de/users/anders/HTSeq) and one of several possible gene model databases. The raw count matrix generated by HTSeq was then processed using the R/Bioconductor package DESeq (http://www-huber.embl.de/users/anders/DESeq) which is used to both normalize the full dataset and analyze differential expression between sample groups. The data was clustered in several ways using the normalized counts of all genes that a total of 10 counts when summed across all samples; 1. Hierarchical cluster with the correlation metric (Dij = 1 - cor(Xi,Xj)) with the Pearson correlation on the normalized log2 expression values. 2. Multidimensional scaling. 3. Principal component analysis. Heatmaps were generated using the heatmap.2 function from the gplots R package. For the Heatmaps the top *100* differentially expressed genes are used. The data plot represents the mean-centered normalized log2 expression of the top 100 significant genes. We ran a gene set analysis using the GSA package with gene sets from the Broads mSigDb. The sets used were: Mouse: c1, c2, c3, c4, c5. Gene ontology analyses were performed using the Database for Annotation, Visualization, and Integrated Discovery (DAVID) Bioinformatics resource (Version 6.8) gene ontology functional annotation tool (http://david.abcc.ncifcrf.gov/tools.jsp) with all NCBI Mus musculus genes as a reference list. KEGG pathway analysis was performed using the KEGG Mapper – Search Pathway function (https://www.genome.jp/kegg/tool/map_pathway2.html). We performed a manual literature search to determine the proportion of significantly changing genes associated with cancer progression and metastasis.

### Accession Numbers

The Gene Expression Omnibus accession number for the RNA-sequencing data reported in this study is GSE148514.

## Acknowledgements

We thank members of the Hadjantonakis and Massagué labs for critical discussions and comments on the manuscript. We also thank members of MSKCC’s Mouse Genetics and Integrated Genomics Operation (IGO) and Bioinformatics Core facilities. All cores are funded by the NCI Cancer Center Support Grant (CCSG, P30 CA08748) and IGO is additionally funded by Cycle for Survival, and the Marie-Josée and Henry R. Kravis Center for Molecular Oncology. SMM is supported by a Wellcome Trust Sir Henry Wellcome postdoctoral fellowship under the supervision of JN and AKH. Work in the Hadjantonakis lab was supported by grants from the NIH (R01HD094868, R01DK084391 and P30CA008748).

## Ethics

Animal experimentation: Animal experimentation: All mice used in this study were maintained in accordance with the guidelines of the Memorial Sloan Kettering Cancer Center (MSKCC) Institutional Animal Care and Use Committee (IACUC) under protocol number 03-12-017 (PI Hadjantonakis).

## Supplemental Figure Legends

**Figure S1. *Rreb1* expression pattern during mouse embryonic development. A.** *Rreb1* expression in different cell types of the early mouse embryo, from published single-cell RNA-sequencing datasets. Left panel: Force-directed layout plot showing relative *Rreb1* expression in cells of E3.5-4.5 pre-implantation and E5.5 early post-implantation embryos from single cell sequencing (sc seq.) data. Plot was generated using data from Nowotschin et al. (Nowotschin et al., 2019). Right panel: Uniform manifold approximation and projection (UMAP) plot, generated using single cell sequencing data from Pijuan-Sala et al. (Pijuan-Sala et al., 2019), showing *Rreb1* expression levels in all the cells at E6.5, 6.75, 7.0 and 7.75. **B.** Schematic diagram showing the original EUCOMM knockout-first (Tm1a) allele (upper panel) and the *Rreb1* null LacZ reporter (Tm1b) allele generated by Cre-mediated recombination of Tm1a (lower panel). *Engrailed 2* splice acceptor (En2 SA), internal ribosome entry side (IRES), human beta actin promoter (hbactP), Neomycin cassette (neo), single polyadenylation sequences (pA), *FRT* sites (green triangles), *loxP* sites (orange triangles). **C.** Wholemount images of E4.5 *Rreb1*^LacZ/+^ reporter blastocysts. **D.** Transverse cryosection through a distal region of an E7.5 *Rreb1*^LacZ/+^ reporter embryos from Figure 1A. Arrowhead indicates expression within the distal anterior epiblast. **E-G.** Wholemount images of *Rreb1*^LacZ/+^ reporter embryos. **H.** Transverse cryosection of the yolk sac of an E10.5 *Rreb1*^LacZ/+^ reporter embryo. **I.** Wholemount images of wild-type embryos following in situ hybridization with sense (control) and antisense probes against *Rreb1*. TE, trophectoderm; ICM, inner cell mass; PrE, primitive endoderm; VE, visceral endoderm; ExE, extraembryonic ectoderm; PS, primitive streak; DE, definitive endoderm; Epi, epiblast; Noto, notochord; Meso, mesoderm; ne, neurectoderm; pcp, prechordal plate; Pr, proximal; Ds, distal; A, anterior; P, posterior; L, left; R, right; ExVE, extraembryonic visceral endoderm; Ch, chorion; Am, amnion; Al, allantois; hf, headfolds; ys, yolk sac; pa i, pharyngeal arch 1; fnp, frontonasal process; lb, limb bud; is, isthmus.

**Figure S2. *Rreb1* mutant embryos exhibit defects at midgestation. A.** Quantification of the proximal to distal length of *Rreb1* wild-type (*Rreb1*^+/+^) and heterozygous (*Rreb1*^+/−^) versus mutant (*Rreb1*^−/−^) littermates at E6.5 (3 litters) and 7.5 (5 litters). Each point represents an individual embryo. Total number of embryos is shown on the graph. Data is shown relative to the average wild-type/heterozygote proximo-distal length of each litter. Bars represent mean and IQR. ** *p* = ≤ 0.005, unpaired t-test. **B.** Brightfield images of wild-type (*Rreb1*^+/+^) and mutant (*Rreb1*^−/−^) littermates at embryonic day (E) 7.75, 8.0 and 9.0. *Rreb1*^−/−^ embryos are smaller than wild-type littermates and do not show stage-appropriate morphological landmarks. **C.** Quantification of relative somite number in E8.5-9.5 *Rreb1* wild-type (*Rreb1*^+/+^) and heterozygous (*Rreb1*^+/−^) versus mutant (*Rreb1*^−/−^) littermates. Each point represents an individual embryo. Data is shown relative to the average somite number of each litter. Separate litters are indicated by different colored points. Bars represent mean and IQR. **E-F.** Transverse cryosections of E9.0 *Rreb1* heterozygous and homozygous mutant, *Afp-*GFP littermates. Boxes mark the regions shown in higher magnification in H. Asterisks mark the open neural tube and gut tube in *Rreb1*^−/−^. Sb, 50 μm. **G.** Confocal optical sections of transverse cryosections from E9.0 embryos in the region of the notochord. From left to right, images show sections from rostral to caudal regions of the anterior embryo. Sb, 20 μm. Pr, proximal; Ds, distal; A, anterior; P, posterior; L, left lateral; R, right; D, dorsal; V, ventral; Am, amnion; Al, allantois; HF, headfolds; ML, midline; n, notochord; nt, neural tube; fg, foregut; ys, yolk sac; pcp, prechordal plate; hb, hindbrain; op, otic pit; ba, branchial arch; fb, forebrain; mb, midbrain.

**Figure S3. *Rreb1*^−/−^ embryos exhibit cardiovascular defects. A.** Uniform manifold approximation and projection (UMAP) plot, generated using single cell sequencing data from Pijuan-Sala et al. (Pijuan-Sala et al., 2019). Left plot shows distinct clusters of cells representing different cell types within the embryo. Adjacent plots show the expression pattern of example genes that were significantly downregulated in *Rreb1*^−/−^ embryos and whose expression is enriched within endoderm tissues. **B.** Graph showing the relative expression level of a panel of endoderm-associated genes from RNA-sequencing of individual *Rreb1*^+/+^ and *Rreb1*^−/−^ embryos that showed no significant difference in expression between genotypes. Each point represents a single embryo. Statistical analysis was performed using an Unpaired *t*-test. Bars represent median and IQR. Expression is shown relative to the mean expression in wild-type embryos. **C.** Diagram illustrating the breeding scheme used to generate Afp-GFP^Tg/+^; *Rreb1*^+/+^ and *Rreb1*^−/−^ embryos. **D.** Confocal MIPs of immunostained embryos *Afp*-GFP; *Rreb1*^+/+^ and *Rreb1*^−/−^ embryos. Arrowheads mark highlight the proximal ExVE that, in contrast to wild-type embryos, shows little to no *Afp*-GFP expression. Sb, 50 μm. **E.** Wholemount images of E10.5 *Rreb1*^LacZ/+^ (heterozygous) and *Rreb1*^LacZ/LacZ^ (mutant) embryos within the yolk sac. Mutant embryos have reduced yolk sac vasculature and blood leaking into the extravascular space (arrowheads). **G.** Brightfield image of two distinct E10.5 *Rreb1*^−/−^ embryos with reduced cranial vasculature (left) and little blood within the fetus (right). Boxes show regions of higher magnification in adjacent panels. **F-H.** Confocal maximum intensity projections showing the cranial and trunk vasculature of E9.5 embryos from Figure 3D. Sb, 50 μm. PECAM-1 marks vasculature. ENDOGLIN marks endothelial cells as well as hematopoietic, mesenchymal and neural stem cells. **H.** Arrowhead marks large blood vessel not observed in wild-type littermate. A, anterior; P, posterior; Pr, proximal; Ds, distal; D, dorsal; V, ventral; ExVE, extraembryonic VE; EmVE, embryonic VE; DE, definitive endoderm.

**Figure S4. Loss of *Rreb1* alters epiblast adherens junction organization. A.** Diagram showing the methodology for quantification of E-CADHERIN protein levels along epiblast cell junctions. Lines were manually drawn along cell junctions and the relative profile of E-CADHERIN immunostaining fluorescence level along the junction was plotted, with the highest value representing 1. We then calculated the coefficient of variation of E-CADHERIN levels for each individual junction. **B.** Representative relative profile of E-CADHERIN levels in arbitrary units (a.u.) at a single *Rreb1*^+/+^ and *Rreb1*^−/−^ epiblast cell junction. **C.** Quantification of the coefficient of variation of E-CADHERIN immunostaining fluorescence levels at epiblast cell junctions. Each point represents a single cell junction. Bars represent mean and IQR. *** *p* = ≤ 0.0005, unpaired t-test. **D.** Confocal maximum intensity projections of transverse cryosections of a lateral region of the epiblast of immunostained *Rreb1*^+/+^ and *Rreb1*^−/−^ embryos. Sb, 10 μm. **E.** Graph showing the relative expression level of a panel of adhesion-associated genes from RNA-sequencing of individual *Rreb1*^+/+^ and *Rreb1*^−/−^ embryos. Each point represents a single embryo. Statistical analysis was performed using an Unpaired *t*-test (\**p*<0.05, \*\**p*<0.005, \*\*\**p*<0.001). Bars represent median and IQR. Expression is shown relative to the mean expression in wild-type embryos. A, anterior; P, posterior; L, left; R, right.

**Figure S5. *Rreb1* mutant embryos have perturbed epithelial architecture. A.** Confocal optical sections showing transverse cryosections in the extraembryonic region of E6.5 embryos. Arrowheads highlight regions where cell layers are abnormally separated from one another. Sb, 25 μm. **B.** Brightfield images of *Rreb1*^+/+^ and *Rreb1*^−/−^ littermates at embryonic day 7.5. Arrowheads highlight the abnormal accumulation of cells at the anterior embryonic-extraembryonic boundary. **C.** Arrows highlight SOX17-expressing definitive endoderm cells within the wings of mesoderm. Sb, 50 μm (A,B) and 25 μm (C). **D.** Confocal sagittal optical sections of immunostained embryos. The *Rreb1*^−/−^ embryo displays abnormal epithelial folding. Sb, 50 μm. **E.** Schematic depicting methodology for angle measurements. We measured the angle of the elongated nuclear axis of epiblast cells relative to the underlying Laminin basement membrane (BM). Sb, 10 μm. **F.** Quantification of the angle between the elongated nuclear axis and the BM of E6.5 epiblast cells. Bars represent median and IQR. Each point represents a single cell. **G.** Quantification of the coefficient of variation (COV) for the nucleus-BM embryo angle in each embryo (individual points). Bars represent mean and IQR. *** *p* = ≤ 0.0005, unpaired t-test. **H.** Confocal optical sections of transverse cryosections in lateral (i) and anterior (ii) regions of E7.5 embryos. Arrowheads highlight regions where cell layers are abnormally separated from one another. Sb, 25 μm. **I.** Confocal optical sections of transverse cryosection of immunostained E7.5 *Rreb1*^−/−^ embryo. Arrowheads highlight a break in apical F-ACTIN through which epiblast cells are protruding. Box indicates region shown at higher magnification. Sb, 25 μm. **J.** Confocal optical sections of transverse cryosections of immunostained E7.5 embryos. In wild-type embryos, epiblast cells divide adjacent to the cavity (arrowheads), maintain apical F-ACTIN and remain within the epithelium. In *Rreb1*^−/−^ embryos, we also observed dividing cells outside of the epithelium (arrowheads), within the amniotic cavity. Sb, 25 μm. **K.** Confocal maximum intensity projections (left) and optical sections of transverse cryosections of immunostained embryos stained for phosphorylated Histone H3 (pHH3), which marks mitotic cells. Sb, 50 μm. **L.** Quantification of proliferation in *Rreb1*^+/+^ and *Rreb1*^−/−^ littermates. We quantified the absolute number of pHH3-positive cells per 10 μm cryosection (left panel) and the % of pHH3 mitotic cells in each germ layer per 10 μm cryosection (right panel) for 3 entire embryos. There was no significant difference (unpaired *t*-test) in proliferation rate between genotypes, other than in the ExE, which is likely a reflection of the low sample number in that region. Each point represents a single dividing cell. Bars represent mean and IQR. **M.** Transverse cryosection of a lateral region of E7.5 epiblasts immunostained for the basal marker, Laminin, and apical marker, ZO-1. *Rreb1*^−/−^ embryos maintain appropriate expression of polarity markers. To note, we observed strong anti-N-CADHERIN and ZO-1 VE fluorescence, which correlates with an apparent difference in the structure of the outer VE layer compared to wild-type embryos. This signal is also observed with other antibodies and likely represents non-specific binding. Sb, 25 μm. **N.** Histogram showing fluorescence levels, in arbitrary units (a.u.), of Laminin and ZO-1 immunostaining measured along the apical-basal axis of a representative region of the epiblast epithelium from image in panel. Pr, proximal; Ds, distal; A, anterior; P, posterior; L, left; R, right; PS, primitive streak; Endo, endoderm; Epi, epiblast; ExVE, extraembryonic visceral endoderm; Meso, mesoderm; ExE, extraembryonic ectoderm.

**Figure S6. Loss of *Rreb1* promotes invasive cell behaviors. A.** Graph showing the expression level in arbitrary units (a.u.) of *Ovol1* and *Cldn7* from RNA-sequencing of individual *Rreb1*^+/+^ and *Rreb1*^−/−^ embryos. Each point represents a single embryo. \*\*\**p*<0.001, unpaired *t*-test. Bars represent median and IQR. **B.** Confocal optical sections showing transverse cryosections through immunostained embryos. Sb, 50 μm. In the *Rreb1*^−/−^ shown in the lower panel, SNAIL is expressed laterally on either side of the posterior epiblast rather than at the posterior pole. Thus, it is unclear whether this expression demarcates a primitive streak-like structure in this case (PS?). **G.** Arrowheads indicate ectopic SNAIL expression in epiblast cells. White lines demarcate a region containing a large cluster of epiblast cells ectopically expressing SNAIL, which exhibit more punctate β-CATENIN localization than in surrounding SNAIL negative epiblast cells. Pr, proximal; Ds, distal; A, anterior; P, posterior; L, left; R, right; Meso, mesoderm; Endo, endoderm; Epi, epiblast; PS, primitive streak. **C.** Confocal optical sections of transverse cryosection of immunostained E7.5 *Rreb1*^−/−^ embryo. Arrowhead marks ectopic cells, in the upper panel, SOX2+ cells leaving the epiblast and traversing the outer endoderm layer and in the lower panel, GATA6+ mesoderm cells traversing the epiblast. Arrow marks SOX2+ debris on the outside of the embryo which may represent dead cells. Sb, 25 μm. **D.** Images highlighting a chain of cells apparently exiting the epiblast and traversing the outer endoderm layer. Chain of cells is artificially colored in orange in lower panel. **E.** Confocal sagittal optical section (upper panel) and maximum intensity project (MIP) (lower panel) of an immunostained E7.5 chimeric embryo containing *Rreb1*^−/−^ ESCs. Arrowheads indicate ectopic SOX2+ cells. Sb, 50 μm. **F.** Quantification of SOX2 protein levels in arbitrary units (a.u.) in normal anterior (aEpi) and posterior (pEpi) Epi cells and SOX2 high (SOX2^HI^) cells in E7.5 *Rreb1^−/−^* chimeric embryos. Data shown relative to mean SOX2 levels within typical aEpi cells. Each point represents a measurement from an individual nucleus (n=696 cells, ***p<0.0001). **G.** Graph showing the proportion of SOX2^HI^ cells localized inside the Epi, at the Epi-VE interface, mesoderm or amniotic cavity in E7.5 *Rreb1^−/−^* chimeric embryos. Data shown as the percentage of the total SOX2^HI^ cells analyzed per embryo in each location. Each point represents scoring for an individual embryo. Total number of cells per location is shown above each bar. For all box plots, top and bottom edges of boxes represent third and first quartiles, respectively (interquartile range, IQR). Middle lines mark the median. Whiskers extend to 1.5 * IQR. **H.** Confocal MIPs of immunostained E8.5 (Sb, 100 μm) and 9.5 (Sb, 200 μm) chimeric embryos containing *Rreb1*^−/−^ ESCs. Arrowheads indicate ectopic SOX2+ cells. **I.** Confocal sagittal optical section of a pre-gastrulation E6.0 chimeric embryo containing *Rreb1*^−/−^ ESCs. Arrowheads mark ectopic SOX2+ cells. mCherry marks ESC progeny. Sb, 25 μm. Boxes show regions displayed at higher magnification. Brackets mark primitive streak. A, anterior; P, posterior; L, left; R, right; Pr, proximal; Ds, distal; Epi, epiblast; Endo, endoderm; Meso; mesoderm; PS, primitive streak; NE, neurectoderm; Am, amnion; Al, allantois; ExE, extraembryonic ectoderm.

**Figure S7. *Rreb1* chimeras display changes in ECM organization. A,C,D.** Confocal optical sections and maximum intensity projections (MIP) of transverse cryosections of immunostained E7.5 chimeric embryos containing *Rreb1*^−/−^ ESCs. Sb, 10 μm. **A.** Cherry fluorescence is a constitutive lineage label marking the progeny of *Rreb1*^−/−^ embryonic stem cells (ESCs) introduced into host embryos. Magenta arrowheads mark ectopic SOX2+ cells derived from *Rreb1*^−/−^ cells. **C.** Arrowhead marks an ectopic break in the basement membrane in a lateral region of the embryo. **D.** Dashed line traces the approximate line of measurement of the Laminin track. Boxes show regions displayed at higher magnification. A, anterior; P, posterior; L, left; R, right; Epi, epiblast; Endo, endoderm; Meso; mesoderm.

**Table S1. List of genes that are differentially expressed between wildtype and Rreb1 mutant embryos.** Differentially-expressed genes were defined as those meeting fold change cutoff log2(2), adjusted p-value cutoff 0.05, and mean coverage of at least 15.

**Table S2. Gene Ontology (GO) analysis of genes significantly upregulated and downregulated in E7.5 Rreb1 mutant embryos.** Gene ontology analyses were performed using the Database for Annotation, Visualization, and Integrated Discovery (DAVID) Bioinformatics resource gene ontology functional annotation tool with all NCBI Mus musculus genes as a reference list.

**Table S3. KEGG pathway analysis of genes significantly upregulated and downregulated in E7.5 Rreb1 mutant embryos.** KEGG pathway analysis was performed using the Database for Annotation, Visualization, and Integrated Discovery (DAVID) Bioinformatics tool.

## Notes

### Competing Interest Statement

The authors have declared no competing interest.

## References

Ahmed, A. R., Griffiths, A. B., Tilby, M. T., Westley, B. R., & May, F. E. (2012). TFF3 is a normal breast epithelial protein and is associated with differentiated phenotype in early breast cancer but predisposes to invasion and metastasis in advanced disease. American Journal of Pathology, 180(3), 904–916. doi:10.1016/j.ajpath.2011.11.022

Aiello, N. M., Maddipati, R., Norgard, R. J., Balli, D., Li, J. Y., Yuan, S., … Stanger, B. Z. (2018). EMT Subtype Influences Epithelial Plasticity and Mode of Cell Migration. Developmental Cell, 45(6), 681–+. doi:10.1016/j.devcel.2018.05.027

Aiello, N. M., & Stanger, B. Z. (2016). Echoes of the embryo: using the developmental biology toolkit to study cancer. Disease Models & Mechanisms, 9(2), 105–114. doi:10.1242/dmm.023184

Ajona, D., Ortiz-Espinosa, S., Pio, R., & Lecanda, F. (2019). Complement in Metastasis: A Comp in the Camp. Frontiers in Immunology, 10. doi:ARTN 669 10.3389/fimmu.2019.00669

Arai, A., Yamamoto, K., & Toyama, J. (1997). Murine cardiac progenitor cells require visceral embryonic endoderm and primitive streak for terminal differentiation. Developmental Dynamics, 210(3), 344–353. doi:10.1002/(Sici)1097-0177(199711)210:3<344::Aid-Aja13>3.0.Co;2-A

Ayollo, D. V., Zhitnyak, I. Y., Vasiliev, J. M., & Gloushankova, N. A. (2009). Rearrangements of the Actin Cytoskeleton and E-Cadherin-Based Adherens Junctions Caused by Neoplasic Transformation Change Cell-Cell Interactions. Plos One, 4(11). doi:ARTN e802710.1371/journal.pone.0008027

Bachir, A. I., Horwitz, A. R., Nelson, W. J., & Bianchini, J. M. (2017). Actin-Based Adhesion Modules Mediate Cell Interactions with the Extracellular Matrix and Neighboring Cells. Cold Spring Harbor Perspectives in Biology, 9(7). doi:ARTN a02323410.1101/cshperspect.a023234

Balmer, S., Nowotschin, S., & Hadjantonakis, A. K. (2016). Notochord morphogenesis in mice: Current understanding & open questions. Dev Dyn, 245(5), 547–557. doi:10.1002/dvdy.24392

Behringer, R., Gertsenstein, M., Vintersten Nagy, K., & Nagy, A. (2014). Manipulating the Mouse Embryo: A Laboratory Manual, Fourth Edition (Vol. 4): Cold Spring Harbor Laboratory Press.

Belaoussoff, M., Farrington, S. M., & Baron, M. H. (1998). Hematopoietic induction and respecification of A-P identity by visceral endoderm signaling in the mouse embryo. Development, 125(24), 5009–5018. Retrieved from <Go to ISI>://WOS:000077976000015

Blockus, H., & Chedotal, A. (2016). Slit-Robo signaling. Development, 143(17), 3037–3044. doi:10.1242/dev.132829

Bradley, A., Anastassiadis, K., Ayadi, A., Battey, J. F., Bell, C., Birling, M. C., … Wurst, W. (2012). The mammalian gene function resource: the international knockout mouse consortium. Mammalian Genome, 23(9-10), 580–586. doi:10.1007/s00335-012-9422-2

Byrd, N., Becker, S., Maye, P., Narasimhaiah, R., St-Jacques, B., Zhang, X. Y., … Grabel, L. (2002). Hedgehog is required for murine yolk sac angiogenesis. Development, 129(2), 361–372. Retrieved from <Go to ISI>://WOS:000173759100009

Cancer Genome Atlas Research Network. Electronic address, a. a. d. h. e., & Cancer Genome Atlas Research, N. (2017). Integrated Genomic Characterization of Pancreatic Ductal Adenocarcinoma. Cancer Cell, 32(2), 185–203 e113. doi:10.1016/j.ccell.2017.07.007

Caswell, D. R., Chuang, C. H., Ma, R. K., Winters, I. P., Snyder, E. L., & Winslow, M. M. (2018). Tumor Suppressor Activity of Selenbp1, a Direct Nkx2-1 Target, in Lung Adenocarcinoma. Mol Cancer Res, 16(11), 1737–1749. doi:10.1158/1541-7786.MCR-18-0392

Chang, H., Dong, T., Ma, X., Zhang, T., Chen, Z., Yang, Z., & Zhang, Y. (2015). Spondin 1 promotes metastatic progression through Fak and Src dependent pathway in human osteosarcoma. Biochem Biophys Res Commun, 464(1), 45–50. doi:10.1016/j.bbrc.2015.05.092

Chen, Q., Yao, Y. T., Xu, H., Chen, Y. B., Gu, M., Cai, Z. K., & Wang, Z. (2016). SPOCK1 promotes tumor growth and metastasis in human prostate cancer. Drug Des Devel Ther, 10, 2311–2321. doi:10.2147/DDDT.S91321

Chen, X. Y., Kojima, S., Borisy, G. G., & Green, K. J. (2003). p120 catenin associates with kinesin and facilitates the transport of cadherin-catenin complexes to intercellular junctions. Journal of Cell Biology, 163(3), 547–557. doi:10.1083/jcb.200305137

Chou, R. H., Wen, H. C., Liang, W. G., Lin, S. C., Yuan, H. W., Wu, C. W., & Chang, W. S. W. (2012). Suppression of the invasion and migration of cancer cells by SERPINB family genes and their derived peptides. Oncology Reports, 27(1), 238–245. doi:10.3892/or.2011.1497

Chung, Y. C., Wei, W. C., Huang, S. H., Shih, C. M., Hsu, C. P., Chang, K. J., & Chao, W. T. (2014). Rab11 regulates E-cadherin expression and induces cell transformation in colorectal carcinoma. Cancer Research, 74(19). doi:10.1158/1538-7445.Am2014-3166

Cofre, J., & Abdelhay, E. (2017). Cancer Is to Embryology as Mutation Is to Genetics: Hypothesis of the Cancer as Embryological Phenomenon. ScientificWorldJournal, 2017, 3578090. doi:10.1155/2017/3578090

Conlon, R. A., & Rossant, J. (1992). Exogenous retinoic acid rapidly induces anterior ectopic expression of murine Hox-2 genes in vivo. Development, 116(2), 357–368. Retrieved from https://www.ncbi.nlm.nih.gov/pubmed/1363087

Damert, A., Miquerol, L., Gertsenstein, M., Risau, W., & Nagy, A. (2002). Insufficient VEGFA activity in yolk sac endoderm compromises haematopoietic and endothelial differentiation. Development, 129(8), 1881–1892. Retrieved from <Go to ISI>://WOS:000175473800008

Deady, L. D., Li, W., & Sun, J. (2017). The zinc-finger transcription factor Hindsight regulates ovulation competency of Drosophila follicles. Elife, 6. doi:10.7554/eLife.29887

Deng, Y. N., Xia, Z. J., Zhang, P., Ejaz, S., & Liang, S. F. (2020). Transcription Factor RREB1: from Target Genes towards Biological Functions. International Journal of Biological Sciences, 16(8), 1463–1473. doi:10.7150/ijbs.40834

Dobin, A., Davis, C. A., Schlesinger, F., Drenkow, J., Zaleski, C., Jha, S., … Gingeras, T. R. (2013). STAR: ultrafast universal RNA-seq aligner. Bioinformatics, 29(1), 15–21. doi:10.1093/bioinformatics/bts635

Dyer, M. A., Farrington, S. M., Mohn, D., Munday, J. R., & Baron, M. H. (2001). Indian hedgehog activates hematopoiesis and vasculogenesis and can respecify prospective neurectodermal cell fate in the mouse embryo. Development, 128(10), 1717–1730. Retrieved from <Go to ISI>://WOS:000169057400002

Elson, E. L. (1988). Cellular Mechanics as an Indicator of Cytoskeletal Structure and Function. Annual Review of Biophysics and Biophysical Chemistry, 17, 397–430. Retrieved from <Go to ISI>://WOS:A1988P532200018

Engstrom, P. G., Steijger, T., Sipos, B., Grant, G. R., Kahles, A., Ratsch, G., … Consortium, R. (2013). Systematic evaluation of spliced alignment programs for RNA-seq data. Nature Methods, 10(12), 1185–+. doi:10.1038/Nmeth.2722

Ferrara, G., & De Vanna, A. C. (2016). Fluorescence In Situ Hybridization for Melanoma Diagnosis: A Review and a Reappraisal. American Journal of Dermatopathology, 38(4), 253–269. doi:10.1097/Dad.0000000000000380

FujimotoNishiyama, A., Ishii, S., Matsuda, S., Inoue, J., & Yamamoto, T. (1997). A novel zinc finger protein, Finb, is a transcriptional activator and localized in nuclear bodies. Gene, 195(2), 267–275. doi:10.1016/S0378-1119(97)00172-8

Gaggioli, C. (2008). Collective invasion of carcinoma cells When the fibroblasts take the lead. Cell Adhesion & Migration, 2(1), 45–47. doi:10.4161/cam.2.1.5705

Gaggioli, C., Hooper, S., Hidalgo-Carcedo, C., Grosse, R., Marshall, J. F., Harrington, K., & Sahai, E. (2007). Fibroblast-led collective invasion of carcinoma cells with differing roles for RhoGTPases in leading and following cells. Nature Cell Biology, 9(12), 1392–U1392. doi:10.1038/ncb1658

Gara, R. K., Kumari, S., Ganju, A., Yallapu, M. M., Jaggi, M., & Chauhan, S. C. (2015). Slit/Robo pathway: a promising therapeutic target for cancer. Drug Discov Today, 20(1), 156–164. doi:10.1016/j.drudis.2014.09.008

Gavara, N., & Chadwick, R. (2016). Relationship between cell stiffness and stress fiber amount, assessed by simultaneous atomic force microscopy and live-cell fluorescence imaging. Biomechanics and Modeling in Mechanobiology, 15(3), 511–523. doi:10.1007/s10237-015-0706-9

George, J. T., Jolly, M. K., Xu, S., Somarelli, J. A., & Levine, H. (2017). Survival Outcomes in Cancer Patients Predicted by a Partial EMT Gene Expression Scoring Metric. Cancer Res, 77(22), 6415–6428. doi:10.1158/0008-5472.CAN-16-3521

Girardi, G., Yarilin, D., Thurman, J. M., Holers, V. M., & Salmon, J. E. (2006). Complement activation induces dysregulation of angiogenic factors and causes fetal rejection and growth restriction. Journal of Experimental Medicine, 203(9), 2165–2175. doi:10.1084/jem.20061022

Gloushankova, N. A., Rubtsova, S. N., & Zhitnyak, I. Y. (2017). Cadherin-mediated cell-cell interactions in normal and cancer cells. Tissue Barriers, 5(3), e1356900. doi:10.1080/21688370.2017.1356900

Goldie, L. C., Nix, M. K., & Hirschi, K. K. (2008). Embryonic vasculogenesis and hematopoietic specification. Organogenesis, 4(4), 257–263. doi:10.4161/org.4.4.7416

Grady, M. E., Composto, R. J., & Eckmann, D. M. (2016). Cell elasticity with altered cytoskeletal architectures across multiple cell types. Journal of the Mechanical Behavior of Biomedical Materials, 61, 197–207. doi:10.1016/j.jmbbm.2016.01.022

Han, M. K. L., & de Rooij, J. (2017). Resolving the cadherin-F-actin connection. Nature Cell Biology, 19(1), 14–16. doi:10.1038/ncb3457

Han, Y. L., Pegoraro, A. F., Li, H., Li, K. F., Yuan, Y., Xu, G. Q., … Guo, M. (2020). Cell swelling, softening and invasion in a three-dimensional breast cancer model. Nature Physics, 16(1), 101–+. doi:10.1038/s41567-019-0680-8

Hui, B. Q., Ji, H., Xu, Y. T., Wang, J., Ma, Z. H., Zhang, C. G., … Zhou, Y. (2019). RREB1-induced upregulation of the lncRNA AGAP2-AS1 regulates the proliferation and migration of pancreatic cancer partly through suppressing ANKRD1 and ANGPTL4. Cell Death & Disease, 10. doi:ARTN 20710.1038/s41419-019-1384-9

Ivanov, A. I., Parkos, C. A., & Nusrat, A. (2010). Cytoskeletal Regulation of Epithelial Barrier Function During Inflammation. American Journal of Pathology, 177(2), 512–524. doi:10.2353/ajpath.2010.100168

Jia, D. Y., Jolly, M. K., Boareto, M., Parsana, P., Mooney, S. M., Pienta, K. J., … Ben-Jacob, E. (2015). OVOL guides the epithelial-hybrid-mesenchymal transition. Oncotarget, 6(17), 15436–15448. doi:DOI 10.18632/oncotarget.3623

Jolly, M. K., Boareto, M., Huang, B., Jia, D. Y., Lu, M. Y., Ben-Jacob, E., … Levine, H. (2015). Implications of the hybrid epithelial/mesenchymal phenotype in metastasis. Frontiers in Oncology, 5. doi:UNSP 15510.3389/fonc.2015.00155

Katsantonis, J., Tosca, A., Koukouritaki, S. B., Theodoropoulos, P. A., Gravanis, A., & Stournaras, C. (1994). Differences in the G/Total Actin Ratio and Microfilament Stability between Normal and Malignant Human Keratinocytes. Cell Biochemistry and Function, 12(4), 267–274. doi:10.1002/cbf.290120407

Kent, O. A., Sandi, M. J., Burston, H. E., Brown, K. R., & Rottapel, R. (2017). An oncogenic KRAS transcription program activates the RHOGEF ARHGEF2 to mediate transformed phenotypes in pancreatic cancer. Oncotarget, 8(3), 4484–4500. doi:10.18632/oncotarget.13152

Ketene, A. N., Roberts, P. C., Shea, A. A., Schmelz, E. M., & Agah, M. (2012). Actin filaments play a primary role for structural integrity and viscoelastic response in cells. Integr Biol (Camb), 4(5), 540–549. doi:10.1039/c2ib00168c

Kim, M., Du, O. Y., Whitney, R. J., Wilk, R., Hu, J., Krause, H. M., … Reed, B. H. (2020). A Functional Analysis of the Drosophila Gene hindsight: Evidence for Positive Regulation of EGFR Signaling. G3-Genes Genomes Genetics, 10(1), 117–127. doi:10.1534/g3.119.400829

Kim, W. K., Kwon, Y., Jang, M., Park, M., Kim, J., Cho, S., … Kim, H. (2019). beta-catenin activation down-regulates cell-cell junction-related genes and induces epithelial-to-mesenchymal transition in colorectal cancers. Sci Rep, 9(1), 18440. doi:10.1038/s41598-019-54890-9

Kofler, N. M., Shawber, C. J., Kangsamaksin, T., Reed, H. O., Galatioto, J., & Kitajewski, J. (2011). Notch signaling in developmental and tumor angiogenesis. Genes Cancer, 2(12), 1106–1116. doi:10.1177/1947601911423030

Kovac, B., Makela, T. P., & Vallenius, T. (2018). Increased alpha-actinin-1 destabilizes E-cadherin-based adhesions and associates with poor prognosis in basal-like breast cancer. Plos One, 13(5), e0196986. doi:10.1371/journal.pone.0196986

Kwon, G. S., Fraser, S. T., Eakin, G. S., Mangano, M., Isern, J., Sahr, K. E., … Baron, M. H. (2006). Tg(Afp-GFP) expression marks primitive and definitive endoderm lineages during mouse development. Dev Dyn, 235(9), 2549–2558. doi:10.1002/dvdy.20843

Kwon, G. S., Viotti, M., & Hadjantonakis, A. K. (2008). The endoderm of the mouse embryo arises by dynamic widespread intercalation of embryonic and extraembryonic lineages. Developmental Cell, 15(4), 509–520. doi:10.1016/j.devcel.2008.07.017

Li, X. M., Wang, J. J., Zhang, C., Lin, C., Zhang, J. M., Zhang, W., … Li, X. N. (2018). Circular RNA circITGA7 inhibits colorectal cancer growth and metastasis by modulating the Ras pathway and upregulating transcription of its host gene ITGA7. Journal of Pathology, 246(2), 166–179. doi:10.1002/path.5125

Liang, O. D., Korff, T., Eckhardt, J., Rifaat, J., Baal, N., Herr, F., … Zygmunt, M. (2004). Oncodevelopmental alpha-fetoprotein acts as a selective proangiogenic factor on endothelial cell from the fetomaternal unit. Journal of Clinical Endocrinology & Metabolism, 89(3), 1415–1422. doi:10.1210/jc.2003-031721

Liang, X., Gomez, G. A., & Yap, A. S. (2015). Current perspectives on cadherin-cytoskeleton interactions and dynamics. Cell Health and Cytoskeleton, 7, 11–24. doi:10.2147/Chc.S76107

Liu, X., Zhou, Z. H., Li, W., Zhang, S. K., Li, J., Zhou, M. J., & Song, J. W. (2019). Heparanase Promotes Tumor Growth and Liver Metastasis of Colorectal Cancer Cells by Activating the p38/MMP1 Axis. Frontiers in Oncology, 9, 216. doi:10.3389/fonc.2019.00216

Mary, S., Charrasse, S., Meriane, M., Comunale, F., Travo, P., Blangy, A., & Gauthier-Rouviere, C. (2002). Biogenesis of N-cadherin-dependent cell-cell contacts in living fibroblasts is a microtubule-dependent kinesin-driven mechanism. Molecular Biology of the Cell, 13(1), 285–301. doi:10.1091/mbc.01-07-0337

Mege, R. M., & Ishiyama, N. (2017). Integration of Cadherin Adhesion and Cytoskeleton at Adherens Junctions. Cold Spring Harbor Perspectives in Biology, 9(5). doi:ARTN a02873810.1101/cshperspect.a028738

Melani, M., Simpson, K. J., Brugge, J. S., & Montell, D. (2008). Regulation of cell adhesion and collective cell migration by hindsight and its human homolog RREB1. Curr Biol, 18(7), 532–537. doi:10.1016/j.cub.2008.03.024

Ming, L., Wilk, R., Reed, B. H., & Lipshitz, H. D. (2013). Drosophila Hindsight and mammalian RREB-1 are evolutionarily conserved DNA-binding transcriptional attenuators. Differentiation, 86(4-5), 159–170. doi:10.1016/j.diff.2013.12.001

Miura, Y., & Wilt, F. H. (1969). Tissue interaction and the formation of the first erythroblasts of the chick embryo. Dev Biol, 19(2), 201–211. doi:10.1016/0012-1606(69)90055-4

Morgani, S. M., Metzger, J. J., Nichols, J., Siggia, E. D., & Hadjantonakis, A. K. (2018). Micropattern differentiation of mouse pluripotent stem cells recapitulates embryo regionalized cell fate patterning. Elife, 7. doi:ARTN e3283910.7554/eLife.32839

Morgani, S. M., Saiz, N., Garg, V., Raina, D., Simon, C. S., Kang, M., … Hadjantonakis, A. K. (2018). A Sprouty4 reporter to monitor FGF/ERK signaling activity in ESCs and mice. Dev Biol, 441(1), 104–126. doi:10.1016/j.ydbio.2018.06.017

Moser, M., & Patterson, C. (2003). Thrombin and vascular development: a sticky subject. Arterioscler Thromb Vasc Biol, 23(6), 922–930. doi:10.1161/01.ATV.0000065390.43710.F2

Mukhopadhyay, N. K., Cinar, B., Mukhopadhyay, L., Lutchman, M., Ferdinand, A. S., Kim, J., … Freeman, M. R. (2007). The zinc finger protein ras-responsive element binding protein-1 is a coregulator of the androgen receptor: implications for the role of the ras pathway in enhancing androgenic signaling in prostate cancer. Molecular Endocrinology, 21(9), 2056–2070. doi:10.1210/me.2006-0503

Nowotschin, S., Setty, M., Kuo, Y. Y., Liu, V., Garg, V., Sharma, R., … Pe'er, D. (2019). The emergent landscape of the mouse gut endoderm at single-cell resolution. Nature, 569(7756), 361–+. doi:10.1038/s41586-019-1127-1

Oliva, C., Molina-Fernandez, C., Maureira, M., Candia, N., Lopez, E., Hassan, B., … Sierralta, J. (2015). Hindsight regulates photoreceptor axon targeting through transcriptional control of jitterbug/Filamin and multiple genes involved in axon guidance in Drosophila. Developmental Neurobiology, 75(9), 1018–1032. doi:10.1002/dneu.22271

Pandey, V., Wu, Z. S., Zhang, M., Li, R., Zhang, J., Zhu, T., & Lobie, P. E. (2014). Trefoil factor 3 promotes metastatic seeding and predicts poor survival outcome of patients with mammary carcinoma. Breast Cancer Res, 16(5), 429. doi:10.1186/s13058-014-0429-3

Pickup, A. T., Lamka, M. L., Sun, Q., Yip, M. L., & Lipshitz, H. D. (2002). Control of photoreceptor cell morphology, planar polarity and epithelial integrity during Drosophila eye development. Development, 129(9), 2247–2258. Retrieved from https://www.ncbi.nlm.nih.gov/pubmed/11959832

Pijuan-Sala, B., Griffiths, J. A., Guibentif, C., Hiscock, T. W., Jawaid, W., Calero-Nieto, F. J., … Gottgens, B. (2019). A single-cell molecular map of mouse gastrulation and early organogenesis. Nature, 566(7745), 490–+. doi:10.1038/s41586-019-0933-9

Pilot, F., Philippe, J. M., Lemmers, C., & Lecuit, T. (2006). Spatial control of actin organization at adherens junctions by the synaptotagmin-like protein Btsz. Nature, 442(7102), 580–584. doi:10.1038/nature04935

Poltavets, V., Kochetkova, M., Pitson, S. M., & Samuel, M. S. (2018). The Role of the Extracellular Matrix and Its Molecular and Cellular Regulators in Cancer Cell Plasticity. Frontiers in Oncology, 8. doi:ARTN 43110.3389/fonc.2018.00431

Reed, B. H., Wilk, R., & Lipshitz, H. D. (2001). Downregulation of Jun kinase signaling in the amnioserosa is essential for dorsal closure of the Drosophila embryo. Current Biology, 11(14), 1098–1108. doi:10.1016/S0960-9822(01)00318-9

Rodriguez, C. I., Buchholz, F., Galloway, J., Sequerra, R., Kasper, J., Ayala, R., … Dymecki, S. M. (2000). High-efficiency deleter mice show that FLPe is an alternative to Cre-loxP. Nature Genetics, 25(2), 139–140. doi:10.1038/75973

Romanienko, P. J., Giacalone, J., Ingenito, J., Wang, Y. J., Isaka, M., Johnson, T., … Mark, W. H. (2016). A vector with a single promoter for in vitro transcription and mammalian cell expression of CRISPR gRNAs. Transgenic Research, 25(2), 259–259. Retrieved from <Go to ISI>://WOS:000371155900158

Saitoh, M. (2018). Involvement of partial EMT in cancer progression. Journal of Biochemistry, 164(4), 257–264. doi:10.1093/jb/mvy047

Sako-Kubota, K., Tanaka, N., Nagae, S., Meng, W. X., & Takeichi, M. (2014). Minus end-directed motor KIFC3 suppresses E-cadherin degradation by recruiting USP47 to adherens junctions. Molecular Biology of the Cell, 25(24), 3851–3860. doi:10.1091/mbc.E14-07-1245

Saxena, K., Srikrishnan, S., Celia-Terrassa, T., & Jolly, M. K. (2020). OVOL1/2: Drivers of Epithelial Differentiation in Development, Disease, and Reprogramming. Cells Tissues Organs, 1–10. doi:10.1159/000511383

Schott, M., de Jel, M. M., Engelmann, J. C., Renner, P., Geissler, E. K., Bosserhoff, A. K., & Kuphal, S. (2018). Selenium-binding protein 1 is down-regulated in malignant melanoma. Oncotarget, 9(12), 10445–10456. doi:10.18632/oncotarget.23853

Sha, Y. T., Haensel, D., Gutierrez, G., Du, H. J., Dai, X., & Nie, Q. (2019). Intermediate cell states in epithelial-to-mesenchymal transition. Physical Biology, 16(2). doi:ARTN 021001 10.1088/1478-3975/aaf928

Stehbens, S. J., Paterson, A. D., Crampton, M. S., Shewan, A. M., Ferguson, C., Akhmanova, A., … Yap, A. S. (2006). Dynamic microtubules regulate the local concentration of E-cadherin at cell-cell contacts. Journal of Cell Science, 119(9), 1801–1811. doi:10.1242/jcs.02903

Su, J., Morgani, S. M., David, C. J., Wang, Q., Er, E. E., Huang, Y. H., … Massague, J. (2020). TGF-beta orchestrates fibrogenic and developmental EMTs via the RAS effector RREB1. Nature, 577(7791), 566–571. doi:10.1038/s41586-019-1897-5

Sun, B., Fang, Y. T., Li, Z. Y., Chen, Z. Y., & Xiang, J. B. (2015). Role of cellular cytoskeleton in epithelial-mesenchymal transition process during cancer progression. Biomedical Reports, 3(5), 603–610. doi:10.3892/br.2015.494

Sun, J. J., & Deng, W. M. (2007). Hindsight mediates the role of Notch in suppressing hedgehog signaling and cell proliferation. Developmental Cell, 12(3), 431–442. doi:10.1016/j.devcel.2007.02.003

Suresh, S. (2007). Nanomedicine - Elastic clues in cancer detection. Nature Nanotechnology, 2(12), 748–749. doi:10.1038/nnano.2007.397

Takahashi, Y., Ohta, T., & Mai, M. (2004). Angiogenesis of AFP producing gastric carcinoma: correlation with frequent liver metastasis and its inhibition by anti-AFP antibody. Oncology Reports, 11(4), 809–813. Retrieved from https://www.ncbi.nlm.nih.gov/pubmed/15010877

Teng, O., Rai, T., Tanaka, Y., Takei, Y., Nakata, T., Hirasawa, M., … Hirokawa, N. (2005). The KIF3 motor transports N-cadherin and organizes the developing neuroepithelium. Cell Structure and Function, 30, 67–67. Retrieved from <Go to ISI>://WOS:000202995600210

Tesar, P. J., Chenoweth, J. G., Brook, F. A., Davies, T. J., Evans, E. P., Mack, D. L., … McKay, R. D. G. (2007). New cell lines from mouse epiblast share defining features with human embryonic stem cells. Nature, 448(7150), 196–U110. doi:10.1038/nature05972

Thiagalingam, A., DeBustros, A., Borges, M., Jasti, R., Compton, D., Diamond, L., … Nelkin, B. D. (1996). RREB-1, a novel zinc finger protein, is involved in the differentiation response to ras in human medullary thyroid carcinomas. Molecular and Cellular Biology, 16(10), 5335–5345. Retrieved from <Go to ISI>://WOS:A1996VH85200011

Vasileva, E., & Citi, S. (2018). The role of microtubules in the regulation of epithelial junctions. Tissue Barriers, 6(3). doi:ARTN e1539596 10.1080/21688370.2018.1539596

Wang, K., Song, Y., Liu, W., Wu, X. H., Zhang, Y. K., Li, S. H., … Yang, C. (2017). The noncoding RNA linc-ADAMTS5 cooperates with RREB1 to protect from intervertebral disc degeneration through inhibiting ADAMTS5 expression. Clinical Science, 131(10), 965–979. doi:10.1042/Cs20160918

Wang, K., Xu, C., Li, W. J., & Ding, L. (2018). Emerging clinical significance of claudin-7 in colorectal cancer: a review. Cancer Management and Research, 10, 3741–3752. doi:10.2147/Cmar.S175383

Wieschaus, E., Nussleinvolhard, C., & Jurgens, G. (1984). Mutations Affecting the Pattern of the Larval Cuticle in Drosophila-Melanogaster .3. Zygotic Loci on the X-Chromosome and 4th Chromosome. Wilhelm Rouxs Archives of Developmental Biology, 193(5), 296–307. doi:10.1007/Bf00848158

Wilk, R., Pickup, A. T., Hamilton, J. K., Reed, B. H., & Lipshitz, H. D. (2004). Dose-sensitive autosomal autonomous and tissue nuclear modifiers identify candidate genes for tissue nonautonomous regulation by the Drosophila zinc-finger protein, hindsight. Genetics, 168(1), 281–300. doi:10.1534/genetics.104.031344

Wilk, R., Reed, B. H., Tepass, U., & Lipshitz, H. D. (2000). The hindsight gene is required for epithelial maintenance and differentiation of the tracheal system in Drosophila. Dev Biol, 219(2), 183–196. doi:10.1006/dbio.2000.9619

Wilt, F. H. (1965). Erythropoiesis in the Chick Embryo: The Role of Endoderm. Science, 147(3665), 1588–1590. doi:10.1126/science.147.3665.1588

Xu, W. W., Mezencev, R., Kim, B., Wang, L. J., McDonald, J., & Sulchek, T. (2012). Cell Stiffness Is a Biomarker of the Metastatic Potential of Ovarian Cancer Cells. Plos One, 7(10). doi:ARTN e4660910.1371/journal.pone.0046609

Yang, J., Antin, P., Berx, G., Blanpain, C., Brabletz, T., Bronner, M., … Temtia, E. I. A. (2020). Guidelines and definitions for research on epithelial-mesenchymal transition. Nature Reviews Molecular Cell Biology, 21(6), 341–352. doi:10.1038/s41580-020-0237-9

Yao, J. J., Zhong, L., Zhong, P. Q., Liu, D. D., Yuan, Z., Liu, J. M., … Liu, B. Z. (2019). RAS-Responsive Element-Binding Protein 1 Blocks the Granulocytic Differentiation of Myeloid Leukemia Cells. Oncology Research, 27(7), 809–818. doi:10.3727/096504018x15451301487729

