## Supplemental Figures for "The transcription factor Rreb1 regulates epithelial architecture and invasiveness in gastrulating mouse embryos"

Morgani et al., Supplemental Figure 1

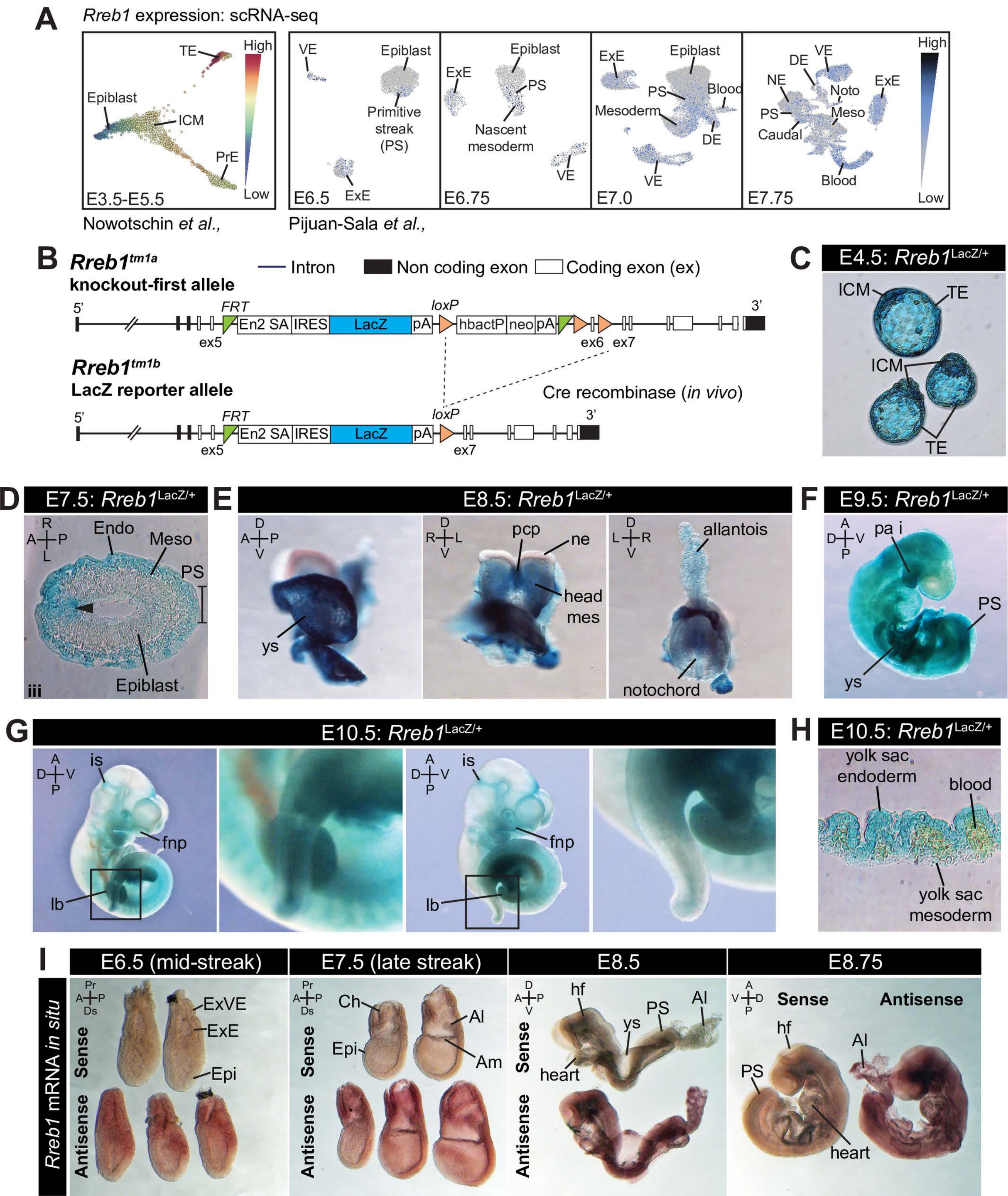

**Morgani et al., Supplemental Figure 2**

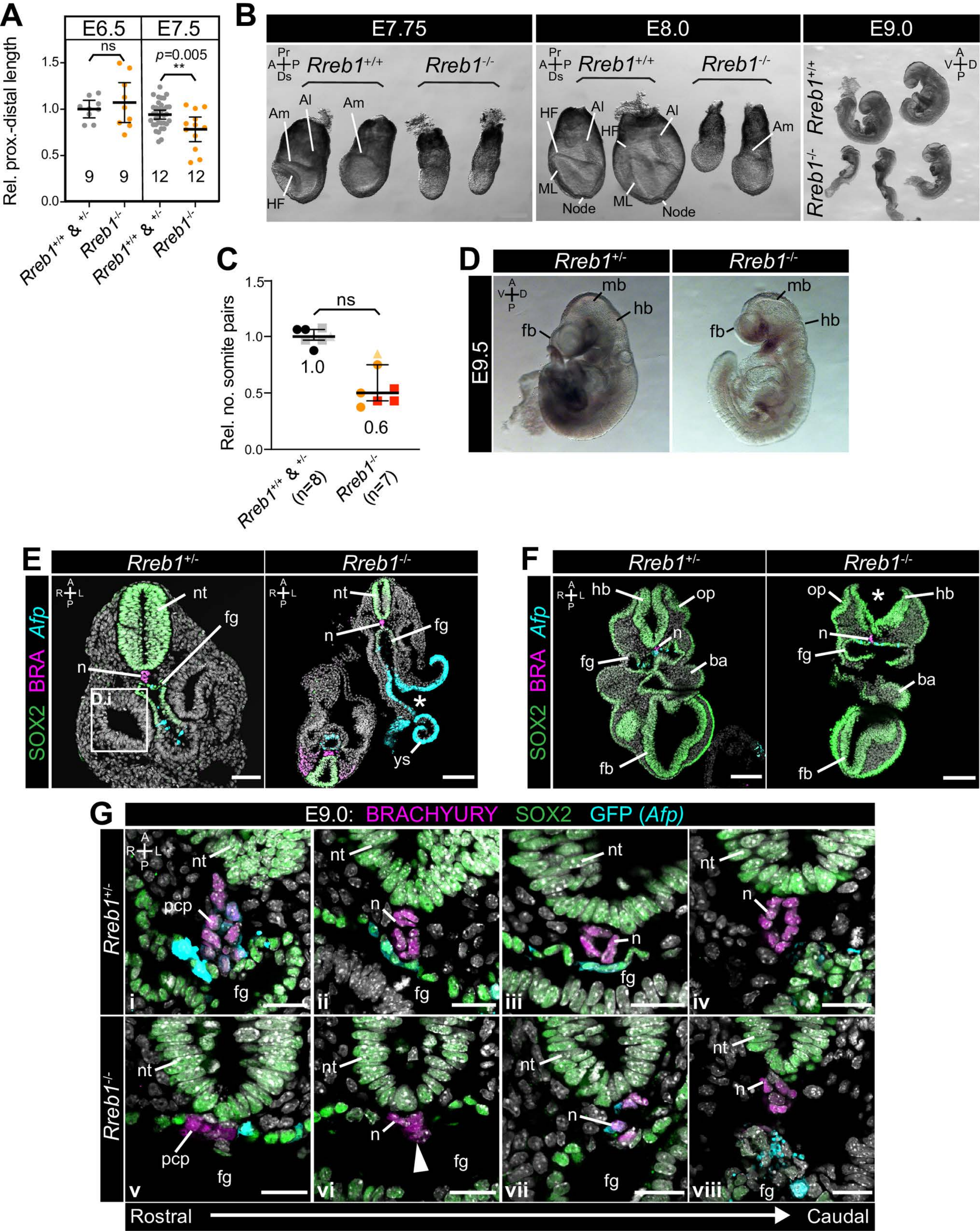

**Morgani et al., Supplemental Figure 3**

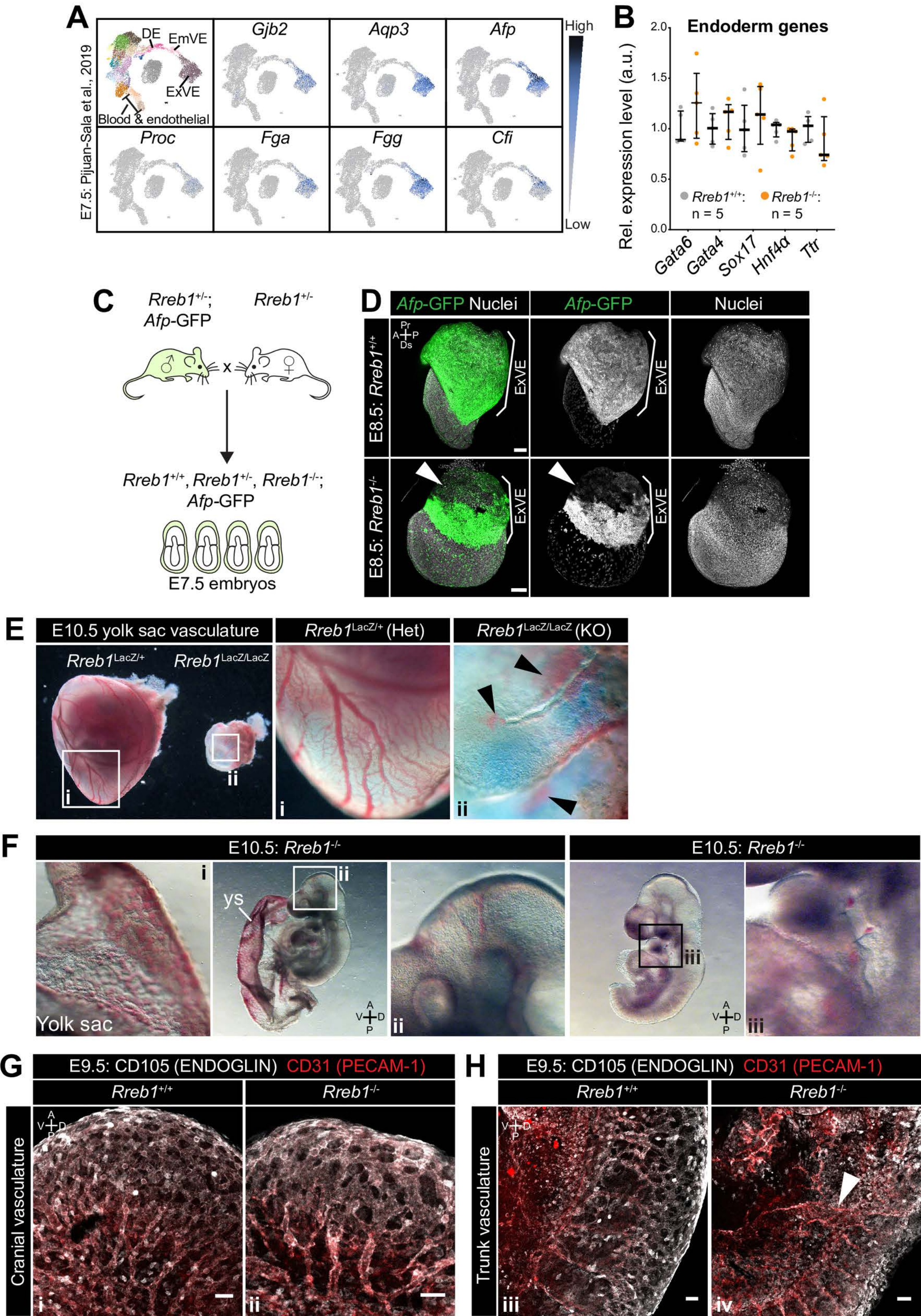

**Morgani et al., Supplemental Figure 4**

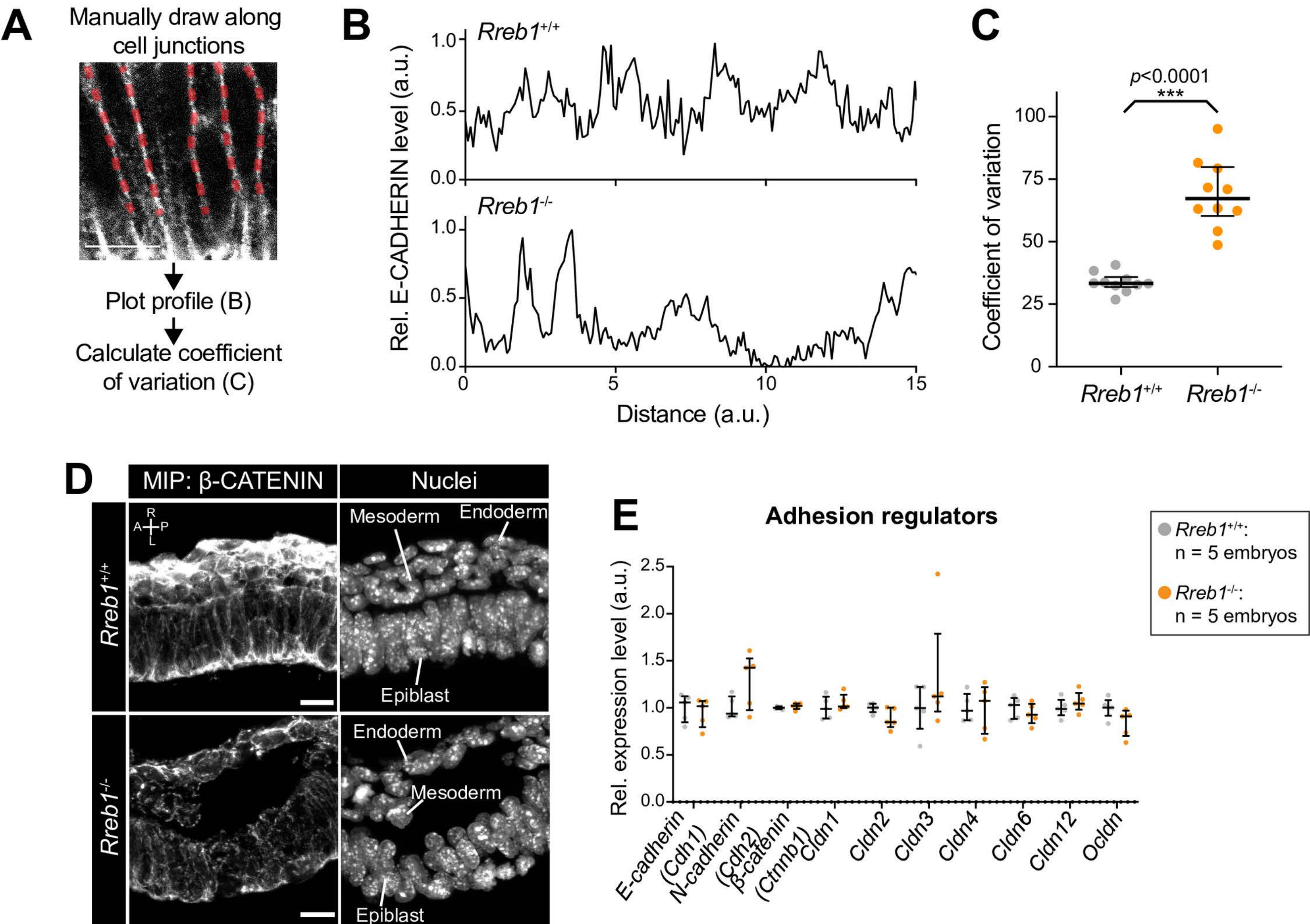

**Morgani et al., Supplemental Figure 5**

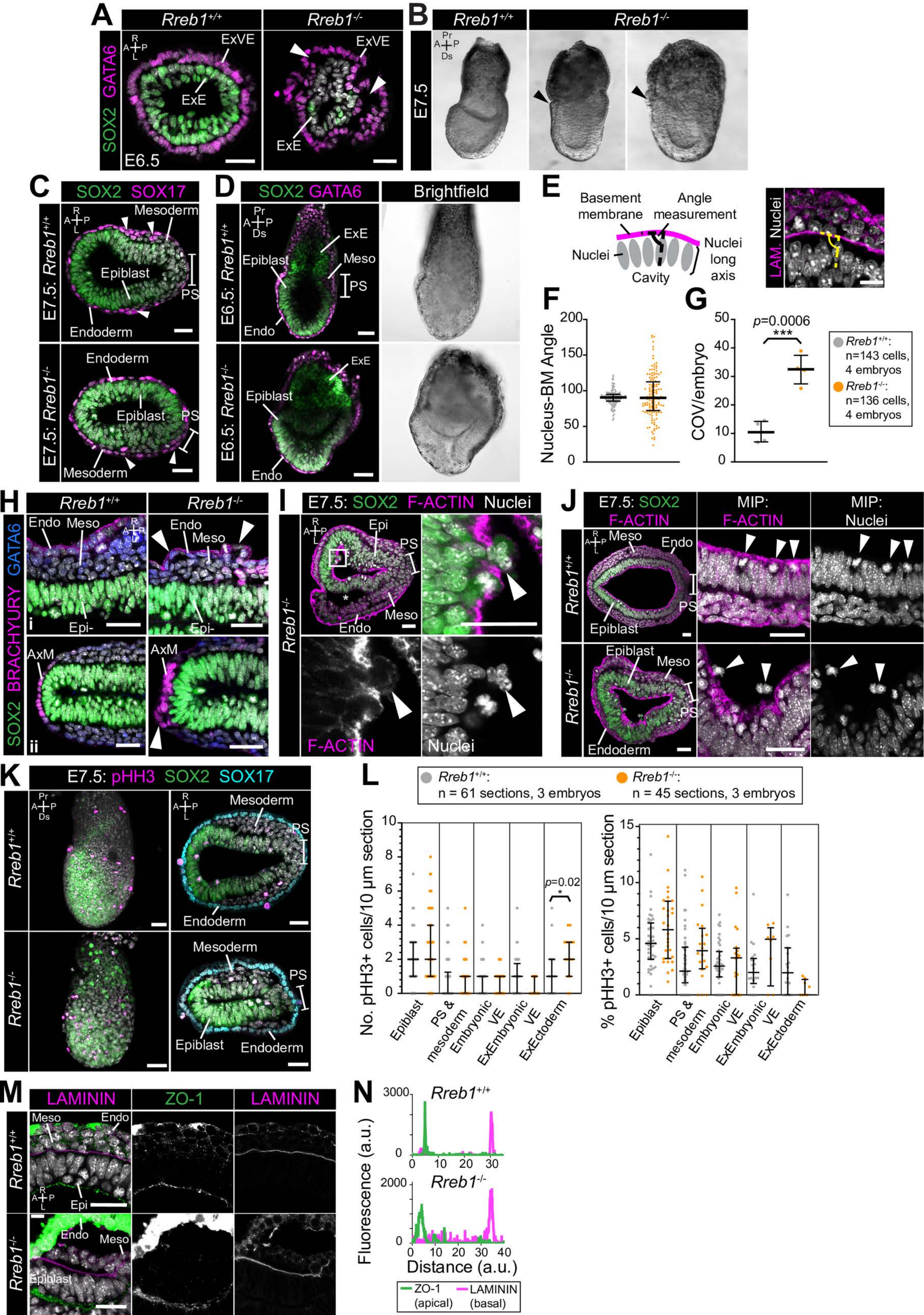

**Morgani et al., Supplemental Figure 6**

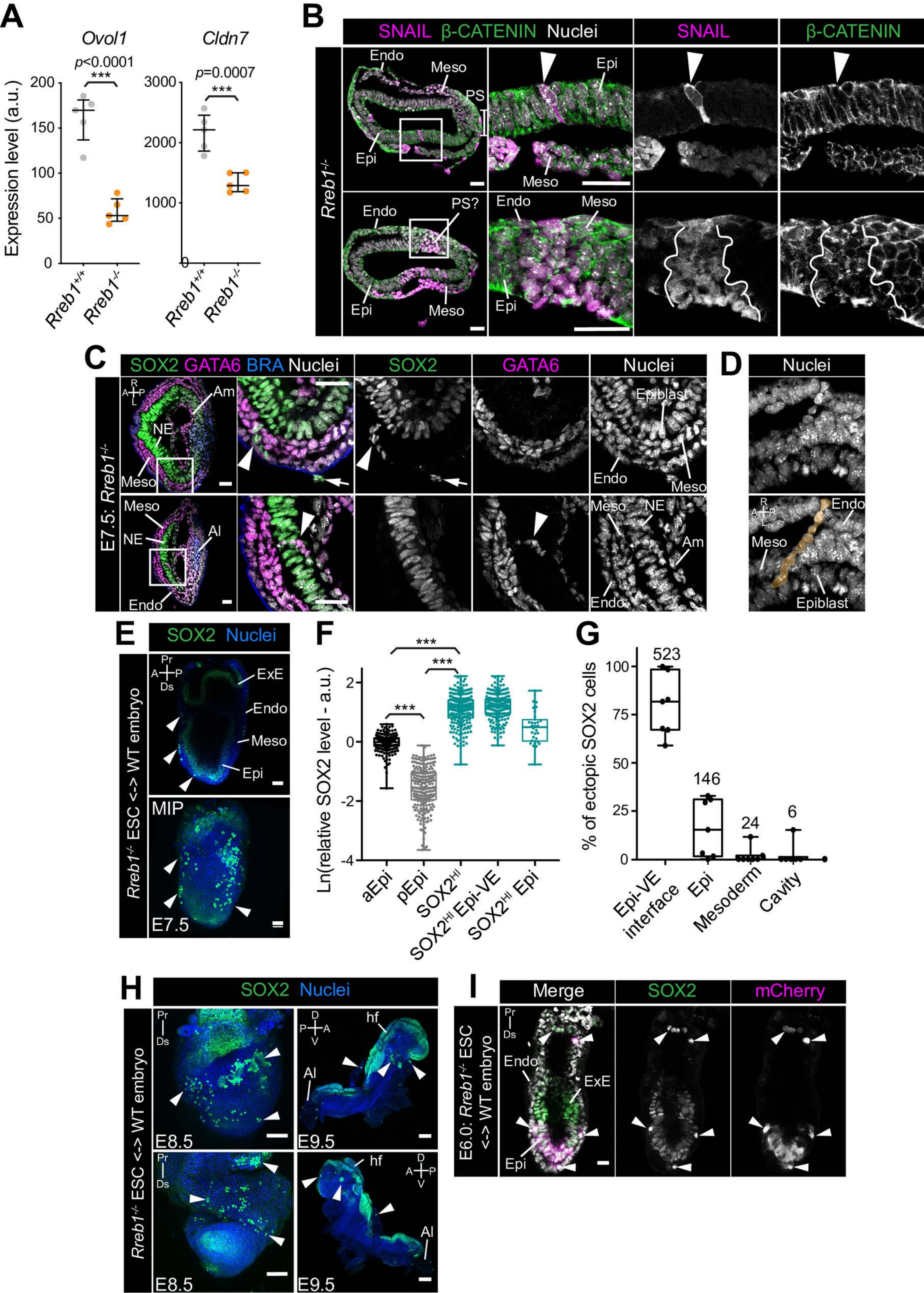

### Morgani et al., Supplemental Figure 7

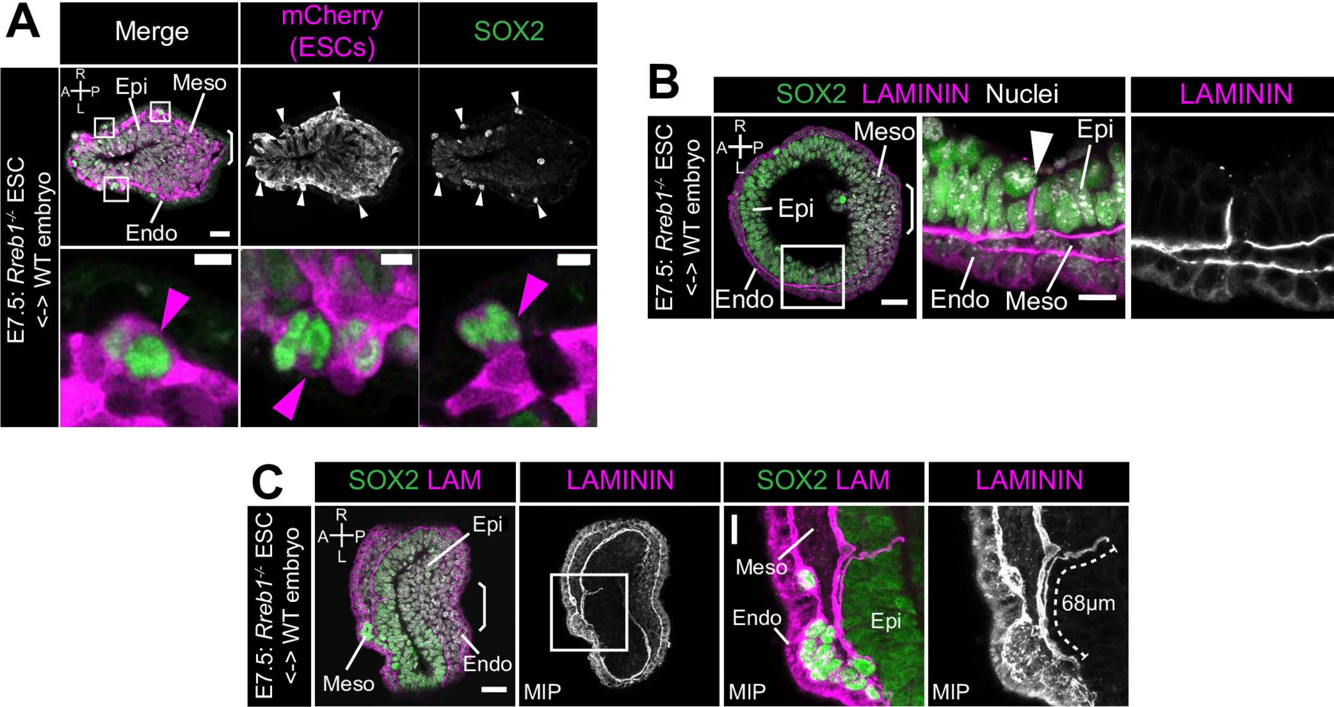
